# Microbial diversity and sulfur cycling in an early earth analogue: From ancient novelty to modern commonality

**DOI:** 10.1101/2021.07.05.451135

**Authors:** C. Ryan Hahn, Ibrahim F. Farag, Chelsea L. Murphy, Mircea Podar, Mostafa S. Elshahed, Noha H. Youssef

## Abstract

Life emerged and diversified in the absence of molecular oxygen. The prevailing anoxia and unique sulfur chemistry in the Paleo-, Meso- and Neoarchean, and early Proterozoic eons may have supported microbial communities that are drastically different than those currently thriving on the earth’s surface. Zodletone spring in southwestern Oklahoma represents a unique habitat where spatial sampling could substitute for geological eons: from the anoxic, surficial light exposed sediments simulating a preoxygenated earth, to overlaid water column where air exposure simulates the relentless oxygen intrusion during the Neo Proterozoic. We document a remarkably diverse microbial community in the anoxic spring sediments, with 340/516 (65.89%) of genomes recovered in a metagenomic survey belonging to 200 bacterial and archaeal families that were either previously undescribed or that exhibit an extremely rare distribution on the current earth. Such diversity is underpinned by the widespread occurrence of sulfite-, thiosulfate, tetrathionate-, and sulfur-reduction, and paucity of sulfate-reduction machineries in these taxa; hence greatly expanding lineages mediating reductive sulfur cycling processes in the tree of life. Analysis of the overlaying water community demonstrated that oxygen intrusion lead to the development of a significantly less diverse community dominated by well-characterized lineages and a prevalence of oxidative sulfur cycling processes. Such transition from ancient novelty to modern commonality underscores the profound impact of the great oxygenation event on the earth’s surficial anoxic community. It also suggests that novel and rare lineages encountered in current anaerobic habitats could represent taxa once thriving in an anoxic earth, but have failed to adapt to earth’s progressive oxygenation.

## Introduction

Sulfur is one of the most abundant elements on earth, exhibiting a wide range of oxidation states (−2 to +6). Microorganisms have evolved a plethora of genes and pathways for exploiting sulfur-redox reactions for energy generation. Reductive processes employ sulfur oxyanions or elemental sulfur as terminal electron acceptors in anaerobic respiratory schemes linked to heterotrophic or autotrophic growth. Oxidative processes, on the other hand, employ sulfides or elemental sulfur as electron donors, powering chemolithotrophic and photosynthetic growth.

Thermodynamic considerations limit reductive sulfur processes to habitats where oxygen is limited. This is reflected in the global distribution of sulfate (SO_4_^2−^), sulfite (SO_3_^2−^), thiosulfate (S_2_O_3_^2−^), tetrathionate (S_4_O_6_^2−^), and elemental sulfur (S^0^) reducing microorganisms (henceforth collectively referred to as SRM) in permanently and seasonally anoxic and hypoxic habitats in marine ^1, 2, 3,^ freshwater ^4,^ terrestrial ^5,^ and subsurface ^6^ ecosystems. Sulfate is highly abundant on the current earth, and hence sulfate-reduction dominates reductive processes in the global sulfur cycle; although reduction and disproportionation of the intermediate sulfur species, e.g. sulfur ^7, 8^, sulfite ^8^, thiosulfate and tetrathionate ^9, 10^ could be significant in localized settings.

The history of earth’s sulfur cycle is a prime example of a geological-biological feedback loop, where the evolution of biological processes is driven by, and dramatically impacts, the earth’s biogeochemistry. The earth’s surface was completely anoxic during the first two billion years of its history, and the availability and speciation of various sulfur species greatly differed from its current values. Sulfate levels were significantly lower when compared to current values in oceanic water (28 mM), with estimates of <200 μM-1mM from the Archean up to the Paleoproterozoic (2.3 Gya) ^11, 12, 13, 14^. On the other hand, intermediate sulfur species appear to have played an important role in shaping the ancient sulfur cycle ^15^. Modeling suggests that mM levels of SO_3_^2−^ were attained in the Archaean anoxic shallow surficial aquifers as a result of dissolution of the volcanic SO_2_ prevailing in aquatic habitats ^12^. Isotopic studies have demonstrated the importance of elemental sulfur, sulfite, and thiosulfate reduction in the Archean^15, 16^.

The evolution of life (3.8 to 4.0 Gy) in the early Archean era and the subsequent evolution of major bacterial and archaeal clades in the Archean and early Proterozoic eons ^17^ occurred within this background of anoxia and characteristic sulfur-chemistry. As such, it has been speculated that organisms using intermediate forms of sulfur were likely more common than sulfate-reducing organisms ^15^. However, while isotopic fractionation, modeling, and microscopic studies could provide clues on prevailing sulfur speciation patterns and prevalent biological processes, the identity of microorganisms mediating such processes is unknown. This is mostly due to constrains on preservation of nucleic acids and other biological macromolecules, with the oldest successful DNA sequenced sample being only 1.2 M years old ^18^. Investigation of the microbial community in modern ecosystems with conditions resembling those prevailing in the ancient earth could provide important clues to the nature, identity, and evolutionary trajectory of microorganisms that thrived under conditions prevailing prior to earth’s oxygenation and the associated changes in the sulfur cycle. In Zodletone spring, a surficial anoxic spring in southwestern Oklahoma, the prevailing conditions are analogous to those predominant on the earth’s surface in the late Archean/early Proterozoic eons At the source of the spring, anoxic, surficial, light-exposed conditions are maintained in the sediments by constant emergence of sulfide-saturated water at the spring source from anoxic underground water formations in the Anadarko basin, along with gaseous hydrocarbons, which occurs in seeps in the general vicinity. These surficial anoxic conditions also support a sulfur chemistry characterized by high levels of sulfide, sulfite, sulfur (soluble polysulfide), thiosulfate, and a low level of sulfate. Further, the sediments at the source of the spring are overlaid by an air-exposed water column, where oxygen intrusion leads to a vertical oxygen gradient (from oxic in the top 1 μm, to hypoxic in the middle, to anoxic in deeper layers overlaying the sediments) that simulates oxygen intrusion into such habitats during the great oxygenation event (Figure S1). As such the spring represents a readily accessible habitat where spatial gradients remarkably correspond to temporal geological transitions. Here, we combined metagenomic, metatranscriptomic, and amplicon-based approaches to characterize the microbial community in Zodletone spring. Our results provide a glimpse of the community mediating the ancient sulfur cycle, significantly expand the overall microbial diversity by the description of a wide range of novel lineages, and greatly increase the number of lineages documented to mediate reductive sulfur processes in the microbial tree of life.

## Results

### Novel phylogenetic diversity in Zodletone sediments

Metagenomic sequencing of the spring sediments yielded 281 Gbp, 79.54% of which assembled into 12 Gbp contigs, with 6.8 Gbp contigs longer than 1Kbp. 1,848 genomes were binned, 683 of which passing quality control criteria, and 516 remaining after dereplication (Table S1). These MAGs represented 64 phyla or candidate phyla (53 bacterial and 11 archaeal), 127 classes, 198 orders, and 300 families (Figure 1a-b). Diversity assessment utilizing small subunit ribosomal protein S3 from assembled contigs (n=2079), as well as a complementary 16S rRNA illumina sequencing effort (n=309,074 amplicons), identified a higher number of taxa (82 phyla and 1679 species in the ribosomal protein S3 dataset, and 69 phyla and 1050 species in 16S rRNA dataset) (Figure S2). Nevertheless, the overall community composition profiles generated from all three approaches were broadly similar (Figure S2), suggesting that the MAG list broadly reflects the sediment microbial community.

**Figure 1.**
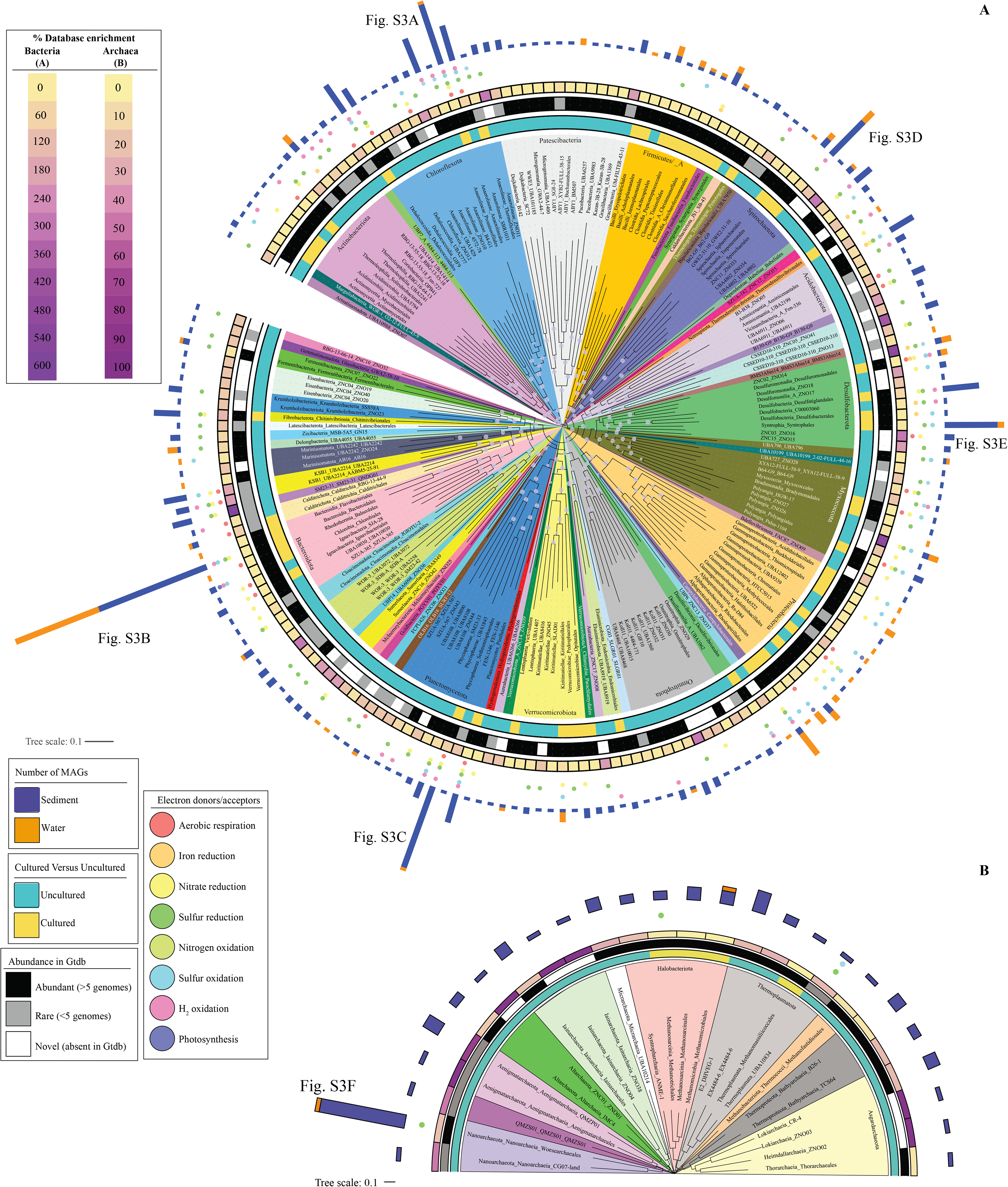
Phylogenomics of the 516 bacterial (A), and 114 archaeal (B) genomes analyzed in this study. The maximum likelihood trees were constructed in FastTree ^84^ based on the concatenated alignments of 120 (bacterial), and 122 (archaeal) housekeeping genes obtained from Gtdb-TK ^83^. The branches represent order-level taxonomy and are color coded by phylum. For phyla with 4 orders or less branches are labeled as Phylum_Class_Order. For phyla with more than 4 orders, the phylum is shown at the base of the colored wedge and the branches are labeled as Class_Order. Lineages staring with ZN depict novel lineages (ZNC = novel class; ZNO = novel order). Bootstrap support values are shown as bubbles for nodes with >70% support. Tracks around the tree represent (from innermost to outermost): cultured status at the order level (cultured versus uncultured), abundance in Gtdb based on the number of available genomes (abundant with more than 5 genomes, rare with 5 genomes or less, and novel with no genomes in Gtdb), percentage database enrichment (calculated as number of genomes belonging to a certain order binned in the current study as a percentage of the number of genomes belonging to the same order in Gtdb), energy conservation capabilities depicted by colored circles (salmon, aerobic respiration; orange, Fe^3+^ respiration; yellow, nitrate/nitrite reduction; dark green, reductive sulfur processes; lime green, nitrogen oxidation; cyan, oxidative sulfur-processes; pink, respiratory hydrogen oxidation; and purple, photosynthesis), and the number of MAGs belonging to each order binned from the sediment (blue bars) and the water (orange bars). For orders with 20 or more genomes, the family-level delineation is shown in Figure 3. These orders are: Anaerolineales (Fig. 3A), Bacteroidales (Fig. 3B), Sedimentisphaerales (Fig. 3C), Spirochaetales (Fig. 3D), Syntrophales (Fig. 3E), and Woesearchaeales (Fig. 3F).

Assessment of novelty and degree of uniqueness of sediment MAGs identified a remarkably high number of previously undescribed lineages (1 phylum, 14 classes, 43 orders, and 97 families), as well as <u>L</u>ineages exhibiting <u>R</u>are global <u>D</u>istribution patterns compared to other present time earth environments (defined as lineages represented by 5 genomes or less in GTDB r95, henceforth LRD) (11 phyla, 24 classes, 45 orders, and 113 families) in the spring. (Figure 1, 2a, b). At the family level, 132 (25.58%), and 208 (40.03%) genomes clustered into 97 novel and 113 LRD families respectively, bringing the proportion of genomes belonging to novel or LRD families in Zodletone sediments to 65.89%. The high level of novelty in the sediment MAGs is reflected in an average RED value of 0.76, a value that is slightly lower than the median RED value for designation of a novel family (0.77) ^19^.

**Figure 2.**
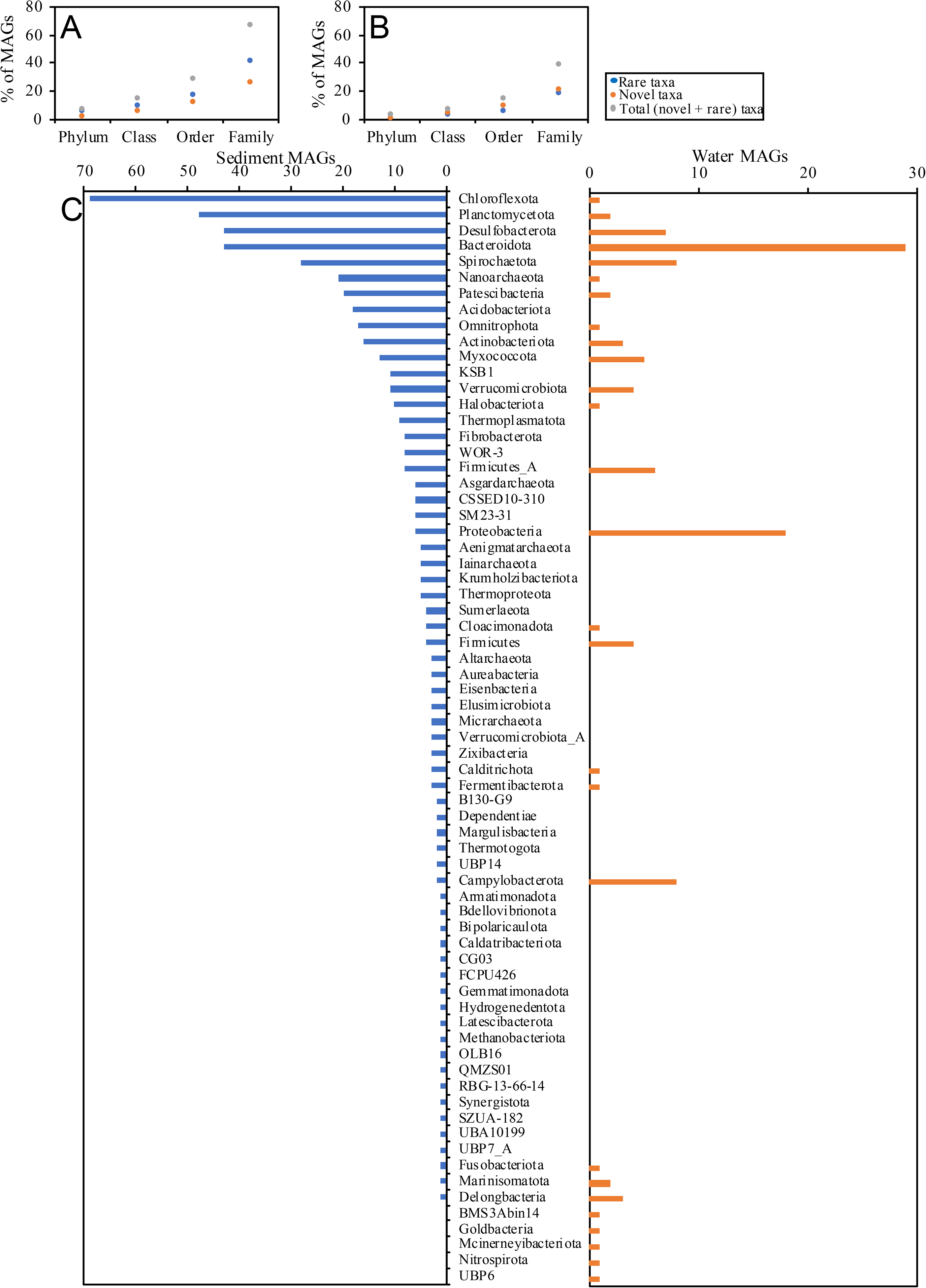
Novelty, rarity, and phylum-level makeup in Zodletone sediment and water communities. Genomes belonging to novel (orange), and LRD (blue) lineages are shown as a percentage of total binned genomes in the sediment (A) and the water (B) communities. The sum of novel and LRD genomes is shown in grey. (C) Phylum-level affiliation for sediment versus water genomes. Number of genomes belonging to each phylum is shown for the sediment (blue bars on the left) and the water (orange bars on the right).

The Chloroflexota (n=69), Planctomycetota (n=47), Bacteroidota (n= 43), Desulfobacterota (n= 43), Spirochaetota (n= 28 genomes), Patescibacteria (n=20 genomes), and the archaeal phylum Nanoarchaeota (n=21) were the most abundant phyla in Zodletone spring sediments, albeit representing only 52.52% of the total number of recovered genomes (S. Text, Figure 1, 2C). An extreme paucity of genomes belonging to the Proteobacteria (6 genomes) and Firmicutes (12 genomes), widely distributed and abundant taxa in current biomes^20, 20,^ and the absence of oxygen-generating Cyanobacteria (0 genomes) were observed (Figure 1, 2C). Within the Chloroflexota, 38/69 genomes belonged to 3 previously undescribed orders, 5 previously undescribed families, and multiple LRD orders and families (Figure 3a). Within the Planctomycetota, 17/47 genomes belonged to 2 previously undescribed orders, 8 previously undescribed families, and multiple LRD orders and families (Figure 3b). Within the Bacteroidota, 27/43 genomes belonged to 1 previously undescribed family, and multiple LRD families (Figure 3c). Within the Spirochaetota, 19/28 genomes belonged to one previously undescribed class, 2 previously undescribed orders, and 9 previously undescribed families, as well as multiple LRD families (Figure 3d). Within the Desulfobacterota (Figure 3e), 35/43 genomes belonged to 3 previously undescribed classes, 10 previously undescribed orders, and 7 previously undescribed families, as well as multiple LRD families. Finally, an extremely diverse community of Patescibacteria (13 different orders, 3 of which belonging to LRD orders, and 14 different families, including 6 previously undescribed and 2 LRD families), and Nanoarchaeota (2 orders and 15 families, including 5 previously undescribed and 10 LRD families) were identified in the spring sediments. A similar pattern of high proportion of previously undescribed and LRD families was identified throughout all other lineages (S. Text, Figure 1a). Therefore, in addition to expanding the number of novel lineages (classes, orders, and families), and greatly enriching available genomes in rare, poorly represented taxa, our results highlight the uniqueness and distinction of the microbial community thriving in Zodletone spring sediments, compared to present earth environments studied so far.

**Figure 3.**
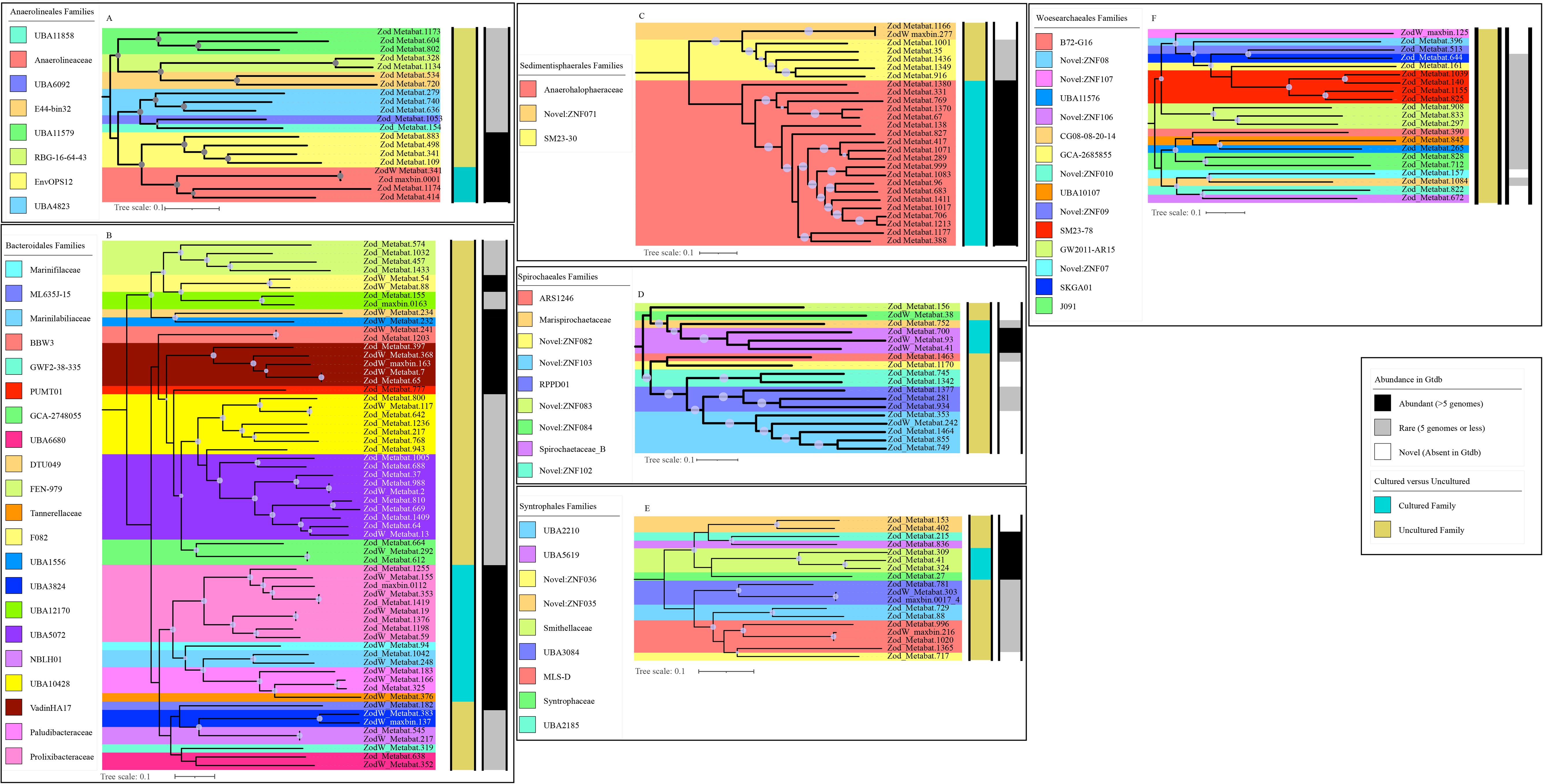
Family-level delineation for orders with 20 or more genomes. The maximum likelihood trees were constructed in FastTree ^84^ based on the concatenated alignments of 120-, and 122-single copy genes, respectively, obtained from Gtdb-TK ^83^. Bootstrap support values are shown as bubbles for nodes with >70% support. Families are color-coded. To the right of the trees, tracks are shown for cultured status at the family level (cultured versus uncultured), and abundance in Gtdb based on the number of available genomes (abundant with more than 5 genomes, rare with 5 genomes or less, and novel with no genomes in Gtdb).

### Oxygen intrusion reduces the proportion of novel and rare lineages in Zodletone spring

Metagenomic sequencing of the oxygen-exposed overlaying water column community yielded 323 Gbp, 80.07% of which assembled into 3.6 Gbp contigs with 3.1 Gbp contigs >1K. 883 genomes were binned, with only 114 remaining after dereplication. Of these, 62 belonged to shared families with the sediment community, and 52 were water specific. Genomes recovered from the water column belonged to a significantly lower number of phyla (n= 27), classes (n=37), orders (n=52), and families (n=79) when compared to the euxinic sediments (Table S1). The community exhibited a much lower level of novelty and rarity at the phylum, class, order, and family levels when compared to the sediment community respectively (Figure 2 a, b). Water-specific genomes (n=52) mostly belonged to well-characterized microbial lineages e.g. families Rhodobacteraceae, and Rhodospirillaceae in the Alphaproteobacteria, families Thiomicrospiraceae, Halothiobacillaceae, Acidithiobacillaceae, Burkholderiaceae, Chromatiaceae, Methylothermaceae in the Gammaproteobacteria, families Sulfurimonadaceae, Sulfurovaceae in phylum Camplylobacterota, and well described families in the phyla Bacteroidota, Desulfobacterota (S. Text, Figure 1, Table S1). Collectively, this demonstrates a pattern where the intrusion of oxygen is associated with a negative impact on previously undescribed and LRD lineages that are prevalent in the sediment, and triggered the propagation of proliferation of cosmopolitan communities within the bacterial tree of life.

#### Reductive sulfur processes dominate Zodletone spring sediment communities

A total of 149 genomes (28.9 % of all genomes), belonging to 32 phyla, 51 classes, 69 orders, and 97 families were involved in at least one reductive sulfur processes (Figure 4, Table S2). By comparison, only 21 sediment genomes (4.06% of all genomes) encoded at least one sulfur oxidation pathway (Figure 4, Table S2). The reductive sulfur-community in the spring exhibited two unique traits: First, a majority of genomes encoding such capacities belonged to novel (47 genomes) or LRD (66 genomes) lineages (Figure 4), and second: sulfite-, polysulfide-, thiosulfate-, and tetrathionate reduction appears to be more prevalent than sulfate-reduction capacities in the sediment genomes.

**Figure 4.**
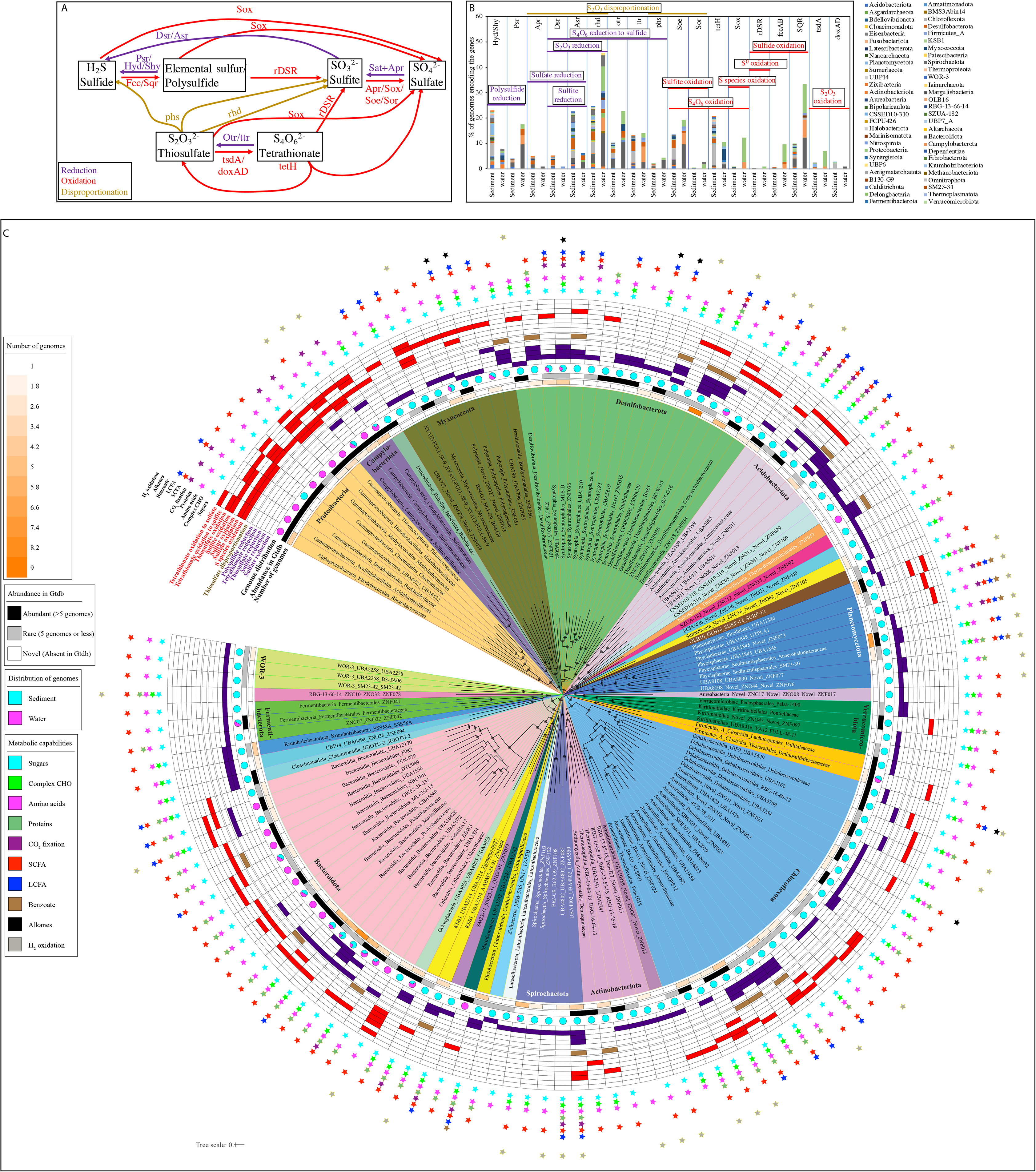
Sulfur cycle in Zodletone spring. (A) Diagram of sulfur transformations predicted to take place in the spring. Different sulfur species are shown in black boxes. Reduction reactions are depicted by purple arrows, oxidation reactions are depicted by red arrows, while disproportionation reactions are depicted by golden brown arrows. The gene(s) names are shown on the arrows. (B) Phylum-level distribution of the S-cycling genes shown at the top of the figure in sediment and water genomes. Processes involving more than one gene are highlighted by horizontal bars and are color coded by reduction (purple), oxidation (red), or disproportionation (golden brown), with the name of the process shown on top of the horizontal bar. (C) Family-level distribution of the genomes involved in S cycling in the spring. The maximum likelihood tree was constructed in FastTree ^84^ based on the concatenated alignments of 120 single-copy genes obtained from Gtdb-TK ^83^. The branches represent family-level taxonomy and are color coded by phylum. For phyla with 2 families or less involved in S cycling, branches are labeled as Phylum_Class_Order_Family. For phyla where 3 or more families are involved in S cycling, the phylum is shown at the base of the colored wedge and the branches are labeled as Class_Order_Family. Lineages staring with ZN depict novel lineages as follows: ZNC, novel class; ZNO, novel order; and ZNF, novel family. Bootstrap support values are shown as bubbles for nodes with >70% support. Tracks around the tree represent (from innermost to outermost): heatmap for the number of genomes in each family, abundance in Gtdb based on the number of available genomes (abundant with more than 5 genomes, rare with 5 genomes or less, and novel with no genomes in Gtdb), pie charts of the breakdown of the number of genomes in the sediment (cyan) versus water (magenta), sulfur reduction pathways (5 tracks in purple), thiosulfate disproportionation pathways (1 track in golden brown), sulfur oxidation pathways (7 tracks in red), and substrates predicted to support growth depicted by colored stars (cyan, sugars; lime green, complex carbohydrates; magenta, amino acids; orange, proteins; purple, CO2 fixation; red, short-chain fatty acids (SCFA); blue, beta oxidation of long-chain fatty acids; brown, anaerobic benzoate/aromatic hydrocarbon degradation; black, anaerobic alkane degradation; and grey, Hydrogen oxidation).

Sulfate-reduction capacity was observed in only 18 sediment genomes (Figure 4, 5a), but exhibited a unique community composition, when compared to well-studied marine and terrestrial habitats ^1, 2, 4, 21^. Sulfate-reduction capacities were observed in mostly previously undescribed or LRD lineages within the Zixibacteria, Acidobacteriota (members of family UBA6911, equivalent to Acidobacteria group 18), Myxococcota, Bacteroidota, Planctomycetota, candidate phylum OLB16 (1 genome), as well as rare and novel lineages within the Desulfobacterota (Figure 4, S3a). Organization of the sulfate reduction genes differed between different phyla, with Myxococcota, Zixibacteria, OLB16, and Acidobacteriota genomes encoding all genes for sulfate activation and reduction, dissimilatory sulfite reduction, as well as energy conservation on one locus, while Desulfobacterota genomes encoded genes for sulfate activation and reduction as well as energy conservation on one locus with genes for dissimilatory sulfite reduction on another, as previously observed in cultured Desulfobacterota ^22, 23, 24^.

**Figure 5.**
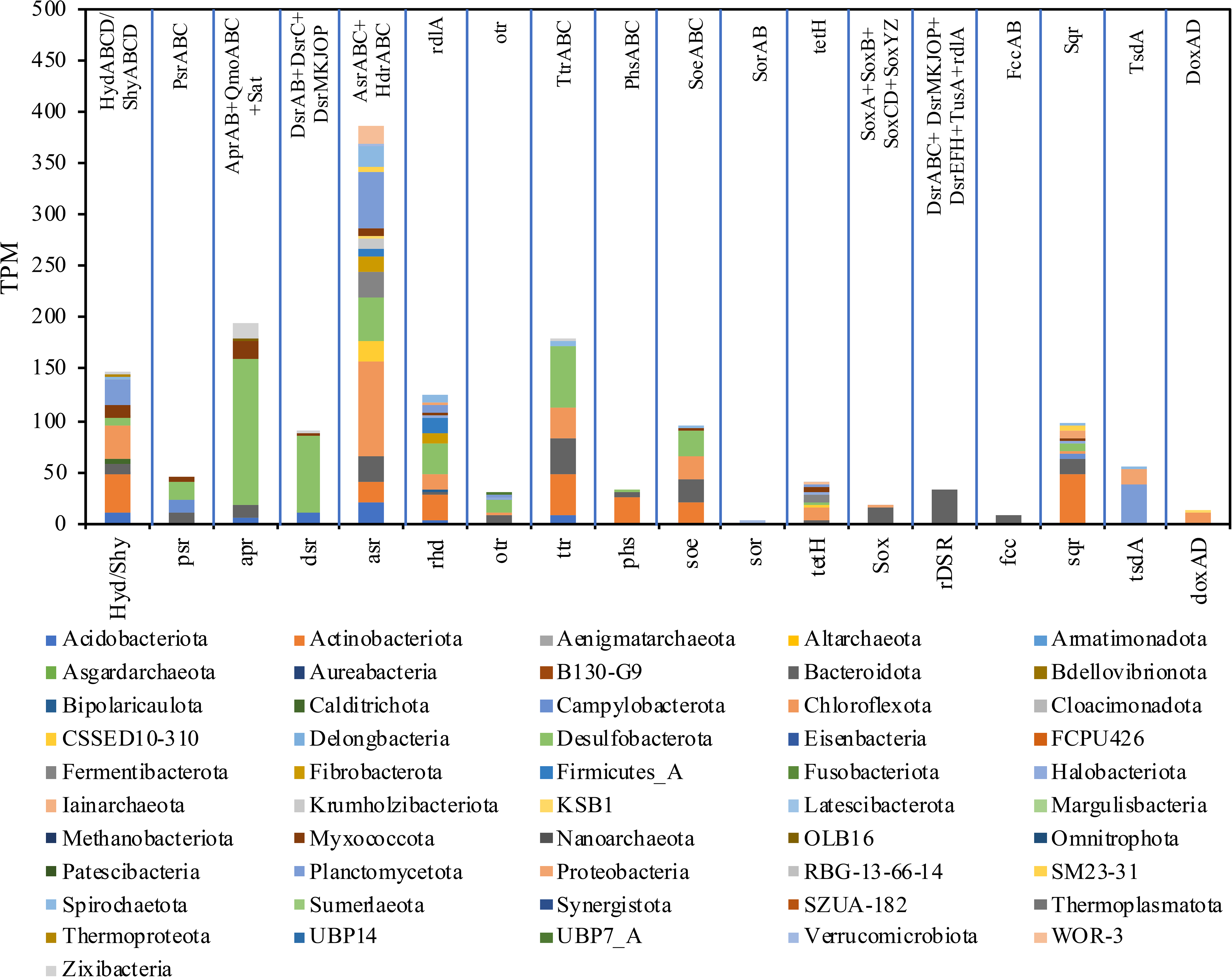
Phylum-level distribution for transcribed sulfur cycling genes in Zodletone spring sediment. RNA-seq reads were pseudo-aligned to the S-cycling genes predicted in Zodletone genomes to detect exact matches using Kallisto ^93^. The transcripts per million are shown on the Y-axis for the gene/ group of genes depicted at the top of the figure.

Sulfite (but not sulfate) reduction via the DsrAB+DsrC+DsrKMJOP system was identified in only 8 genomes belonging to 7 families within the phyla Planctomycetes, Chloroflexota, Spirochaetota, and Desulfobacterota (Figure 4, S3b). Gene organization of the *dsr* locus in the above 9 genomes differed between different phyla, with Chloroflexota, and Spirochaetota genomes encoding all genes for dissimilatory sulfite reduction (dsrABC plus dsrKMJOP) on one locus, while Planctomycetota and Desulfobacterota genomes showed a split *dsr* locus with dsrABC on one locus and dsrMKJOP on another.

On the other hand, sulfite-reduction capacity within Zodletone spring sediment solely via the Asr/Hdr system was rampant, being encountered in 104 genomes belonging to 28 phyla, 43 (8 novel and 9 LRD) classes, 56 (18 novel, and 12 LRD) orders, and 72 (31 novel and 25 LRD) families (Figure 4, S3b), with a gene organization of the *asr* locus adjacent to the *hdr* locus in the majority of genomes. Asr-encoding genomes in the sediments included members of predominantly previously undescribed and LRD lineages within the Chloroflexota, Desulfobacterota. Planctomycetota, and Bacteroidota. The capacity was also rampant in the yet-uncultured bacterial phyla, many of which have fairly limited distribution on the current earth, e.g. the candidate phyla CSSED10-310, FCPU426, RBG-13-66-14, SM23-31, SZUA-182, UBP14, Aureabacteria, Sumerlaeota. Interestingly, all genomes belonging to the novel phylum Krumholzibacteriota, recently described from the spring sediment ^25,^ encoded complete anaerobic sulfite reductase systems. Zodletone dissimilatory sulfite reductase (Figure S3a) and the anaerobic sulfite reductase (Figure S3b) sequences clustered with reference sequences from the same phylum, generally showing no evidence of LGT (S. text). Sulfur (polysulfide) reduction capacities were observed in twenty Zodletone sediment genomes that encoded *psrABC* genes (Figure 4, S3c). These genomes belonged mostly to previously undescribed and LRD families within the phyla Bacteroidota, Desulfobacterota, Myxococota, Acidobacterota, Chloroflexota, and Campylobacterota (Figure 4), In addition, representatives of the cytolpasmic sulfurhydrogenase I (HydABCD system), and/or II (ShyABCD system) were identified in 119 Zodletone sediment genomes (Figure 4). However, as described above, direct involvement of these enzymes in an ETS-associated respiration is not yet clear.

#### Thiosulfate disproportionation and reduction

The quinone-dependent membrane-bound molybdopterin-containing thiosulfate reductase PhsABC was encoded in 11 genomes belonging to 6 phyla, with Bacteroidota representing the major phsABC-encoding phylum (4 genomes). Within these genomes, only two (a Chloroflexota family UBA6092 genome, and a Desulfatiglandales family HGW15 genome) also encoded a dissimilatory sulfite reductase (the Asr system) akin to the Gammaproteobacteria thiosulfate disproportionating pure culture members, where the final products of thiosulfate disproportionation are expected to be only hydrogen sulfide (Figure 4). On the other hand, 5 of the 11 phsABC-encoding Zodletone genomes also encoded the sulfite dehydrogenase SoeABC system, akin to Desulfobacterota and Firmicutes pure culture members, where the final products of thiosulfate disproportionation are expected to be both hydrogen sulfide and sulfate. (Figure 4).

In addition to the phsABC system, 14 Zodletone genomes belonging to 6 phyla (Desulfobacterota,Acidobacteriota, Chloroflexota, Bacteroidota, Spirochaetota, and Myxococcota) encoded a rhodanase-like enzyme [EC: 2.8.1.1 or EC: 2.8.1.3] for thiosulfate disproportionation, as well as enzymes for both sulfite oxidation (by means of reversal of sulfate reduction via Sat+AprAB, or the sulfite dehydrogenase SoeABC), and sulfite reduction (via the dissimilatory sulfite reductases Dsr or Asr), where the final products of thiosulfate disproportionation are expected to be both hydrogen sulfide and sulfate (Figure 4).

#### Tetrathionate reduction

Seventy-three Zodletone sediment genomes encoded the octaheme tetrathionate reductase (OTR) enzyme. These genomes belonged to 14 phyla with major contribution from Bacteroidota (30 genomes), Chloroflexota (10 genomes), and Desulfobacterota (10 genomes). In addition to Otr, 68 Zodletone genomes encoded the Ttr enzyme system. These genomes belonged to 14 phyla with major contribution from Chloroflexota (22 genomes), and Desulfobacterota (20 genomes). As shown previously in *Salmonella typhimurium* ^26,^ in presence of means for thiosulfate disproportionation/reduction and sulfite reduction, the thiosulfate produced as a result of tetrathionate reduction could be further reduced to sulfide. Out of the 105 sediment genomes encoding the Otr, and/or Ttr enzymes, only 12 genomes also encoded thiosulfate and sulfite reduction enzymes. These genomes belonged to the phyla Acidobacteriota, Chloroflexota, Desulfobacterota, Myxococcota, and Spirochaetota.

#### Substrates supporting sulfidogenic capacities at Zodletone spring

Within lineages mediating reductive sulfur processes in Zodletone sediments (n=98), a wide range of substrates supporting sulfidogenesis were identified (Table S3, Figure 4). These included hexoses (26-87% of sulfidogenic lineages), pentoses (30-41% of sulfidogenic lineages), amino acids and peptides (39% of lineages), short chain fatty acids, e.g. lactate, propionate, butyrate, and acetate (22-73% of lineages), long chain fatty acids (29% of lineages), aromatic hydrocarbons (3% of lineages), and short chain alkanes (6% of lineages). Autotrophic capacities with hydrogen as the electron donor were identified in 28% of sulfidogenic lineages.

### Transcriptomic analysis

Transcriptional expression of genes involved in S-species reduction/disproportionation was analyzed in the spring sediments (Figure 5). All S-species reduction/ disproportionation genes discussed above were identified and mapped to 51 distinct phyla. Total transcription levels of the Asr system were 4-times higher than the Dsr system, consistent with the higher number of Zodletone sediment genomes encoding the Asr system compared to the Dsr system. Transcription of Asr system genes were mapped to the phyla Chloroflexota, Planctomycetota, Desulfobacterota, Bacteroidota, Fermentibacterota, Acidobacteriota, CSSED10-310, Actinobacteriota, Spirochaetota, WOR-3, and Fibrobacterota; while DSR genes to Desulfobacterota, Acidobacteriota, Zixibacteria, and Myxococcota (Dsr) (Figure 5). Sulfate reduction genes (Sat, AprAB, and QmoABC) were also transcribed with major contribution from Desulfobacterota, Myxococcota, Zixibacteria, and Acidobacteriota. Transcription of the thiosulfate disproportionating rhodanese-like enzyme [EC: 2.8.1.1 or EC: 2.8.1.3], thiosulfate reductase *phsABC*, tetrathionate reduction genes *ttrABC*, octaheme tetrathionate reductase *otr*, and *psrABC* for polysulfide reduction, and cytoplasmic sulfurhydrogenases I and II (*hyd/shy* systems) was also identified (Figure 5, S. text for detailed contributions of taxa).

### Oxidative sulfur processes dominate Zodletone water community

Reductive sulfur-processes were extremely sparse in the water community (S. text). In contrast oxidative sulfur processes dominated the water community, with pathways encoding sulfide, sulfur, thiosulfate, tetrathionate, and/or sulfite oxidation to sulfate present in 59/114 (51.8%) of water genomes, belonging to 13 phyla, 16 classes, 25 orders, and 43 families, respectively. The oxidative sulfur community in the water belonged to mostly well-characterized lineages (Table S3, Figure 4), with only 8 and 10 genomes involved in oxidative sulfur processes belonged to previously undescribed, and LDR families, respectively. A complete SOX system, putatively mediating oxidation of a wide range of reduced sulfur-species to sulfate was encoded in genomes belonging to well-characterized families within the Proteobacteria (11 genomes in Acidithiobacillaceae, Burkholderiaceae, Halothiobacillaceae, Rhodobacteraceae, and Thiomicrospiraceae) and Campylobacterota (3 genomes in the family Sulfurimonadaceae) (Figure 4). The capacity for sulfide oxidation to sulfur (sulfide dehydrogenase and/or the sulfide:quinone oxidoreductase Sqr) was encoded in thirty nine water genomes within the phyla Bacteroidota, Proteobacteria, Campylobacterota, Chloroflexi, Desulfobacterota, Marinisomatota, and Nitrospirota (S. Text, Figure 4). Only two of the above thirty-nine genomes (a Proteobacteria genome and a Nitrospirota genome) encoded the capacity to further oxidize the sulfur/polysulfide to sulfite via the reversal of the Dsr system, the sulfite dehydrogenase (quinone) SoeABC, or the sulfite dehydrogenase (cytochrome) SorAB system was encountered in 1, 22, and 3 genomes respectively, belonging to well characterized families within Bacteroidetes, Proeobacteria, and Campilobacterota (S. text, Figure S4). Finally, for thiosulfate oxidation, eight water genomes (Proteobacteria and Flavobactereaceae) encoded thiosulfate to tetrathionate oxidation capacities via either the thiosulfate dehydrogenase *tsdA* [EC: 1.8.2.2] or *doxAD* [EC: 1.8.5.2]. Two of these 8 genomes (1 Rhodobacteraceae, and 1 Halothiobacillaceae genomes) also encoded tetrathionate hydrolase (*tetH*) ^27^ known to cleave tetrathionate to thiosulfate, sulfur, and sulfate. Simultaneous identification of the SOX system and both forms of sulfide dehydrogenase (fccAB and Sqr) imply that these two genomes encode the capacity for complete thiosulfate oxidation to sulfate.

## Discussion

The microbial community in Zodletone spring sediments exhibited a high level of phylogenetic diversity, novelty, and rarity (Figure 1–3). Conversely, representatives of lineages that predominate in most present earth environments, e.g. Proteobacteria, Firmicutes, and Cyanobacteria were absent or extremely sparse within the sediments. The community in the spring sediments was also characterized by a high proportion of SRM, and the prevalence of lineages mediating sulfur cycle intermediates (sulfite, thiosulfate, tetrathionate, and elemental sulfur). Many of the organisms mediating reductive sulfur-cycling processes belonged to novel and LRD lineages, hence greatly expanding the range of SRM within the tree of life (Figure 4).

What drives the assembly, propagation, and maintenance of such a diverse, novel, and distinct community in the spring sediments? The high level of diversity, novelty, and rarity within Zodletone spring sediment SRM community could be attributed to two main factors. First, the availability of a wide range of sulfur cycle intermediates in concentrations much higher than sulfate, in contrast to sulfate-predominance in current ecosystems ^2^. Such pattern selects for a more diverse community of SRM in the spring, when compared to predominantly sulfate-driven marine and freshwater ecosystems (Figure 4). Second, additional factors usually constraining SRM growth in several habitats, e.g. diel or seasonal intrusion of oxygen, Fe and NO_3_ ^1, 28, 29, 30,^ recalcitrance of available substrates ^6, 31, 32,^ temperature ^33, 34,^ pH ^21, 35, 36,^ salinity ^37,^ and pressure extremes ^31, 38,^ or combinations thereof, are absent in the spring. Therefore, while the reductive global sulfur cycle appears to be dominated by a few sulfate-reducing lineages within the Desulfobacterota, and to a lesser extent the Firmicutes, as well as Thermodesulfobacteria and *Archaeoglobus* in high temperature habitat, the SRM community in the spring is extremely diverse, encompassing a wide range of previously undescribed and LRD lineages (Figure 4, 5, S4-S6).

Sulfate-reducing organisms are the most prevalent component of the reductive sulfur cycle in most marine and aquatic ecosystems. Aspects of the ecology ^2,^ physiology ^39,^ and biochemistry ^40, 41, 42^ of dissimilatory sulfate reduction have been extensively investigated ^43^. While not the most prevalent process, the sulfate-reducing community in Zodletone spring sediment exhibited a unique composition, with members of the Zixibacteria, Acidobacteriota, Myxococcota, Bacteroidota, Planctomycetota, and candidate phylum OLB16 constituting the major players, as well as rare and novel lineages within the Desulfobacterota (Desulfatiglandales, and Order C00003060), with scarce representation of canonical Desulfobacterota sulfate reducers (1 genome). While the identification of the dissimilatory sulfate reducing machinery in some of these lineages (e.g. Zixibacteria, Acidobacteriota, and Planctomycetota) has been shown before ^44, 45, 46,^ these members rarely appear to be the dominant players in a single ecosystem.

Compared to sulfate-reduction, the ecology and diversity of microbial dissimilatory sulfite reduction has not been extensively studied. The biochemistry of the process has been examined in sulfate-reducers, when grown on sulfite ^43,^ as well as in a few other dedicated sulfite reducers, e.g. *Desulfitobacterium* ^47,^ *Salmonella* ^48,^ *Shewanella* ^49,^ and *Wolinella* ^50^. A recent study suggested the importance and ancient nature of sulfite reduction in an extreme thermophilic environment in a limited diversity biofilm ^51^. We document a plethora of microorganisms within the phyla Planctomycetes, Chloroflexota, Spirochaetota, and Desulfobacterota encoding the dissimilatory sulfite reductase DSR, as well as 72 additional families (31 novel and 25 LRD) encoding the anaerobic sulfite reductase. These organisms greatly expand the known sulfite reduction capacity within the domain Bacteria. Further, the novelty or rarity of some of these families is a reflection of the dearth of present-time habitats that could support this mode of metabolism, once predominant on ancient earth.

The bulk of knowledge on thiosulfate reduction and or disproportionation comes from studies in pure cultures, e.g. members of the Desulfobulbaceae (e.g. *Desulfocapsa*) ^52, 53^ and the genera *Desulfovibrio* and *Desulfomonile* ^54, 55, 56^ within Desulfobacterota, the gammaproteobacterium *Pantoea agglomerans* ^57,^ members of Thermodesulfobacteria ^58^ and Firmicutes ^59^. Radioisotope tracing of different sulfur atoms showed a significant contribution of thiosulfate disproportionation to the sulfur cycle in marine ^60,^ as well as freshwater sediments ^61^. However, the lack of a marker gene for the process hinders ecological culture-independent studies. Similar to sulfite, the high levels, and constant generation of thiosulfate in Zodletone sediments sustains a highly diverse thiosulfate reducing (thiosulfate reductase plus a sulfite reduction complex), or disproportionating (thiosulfate reductase plus both sulfite reduction and sulfite oxidation systems) community with major contribution from novel or rare families in the Acidobacteriota, Chloroflexota, Desulfobacterota, KSB1, Myxococcota, and Spirochaetota. Finally, the extremely high levels of zero valent sulfur, available as soluble polysulfide, result in enriching the community with a plethora of polysulfide-reducing organisms.

As described above, the prevailing conditions in Zodletone spring (anaerobic, surficial, light-exposed, sulfidic, with abundance of S cycle intermediates) remarkably mimic the conditions that were prevalent in the late Archean and early Proterozoic eons, when the evolution of major bacterial and archaeal clades (based on recently refined timing estimates from ^62^), with the notable exception of Proteobacteria and Cyanobacteria) occurring prior to oxygen evolution. This study infers that the microbial communities presumably thriving in surficial earth were extremely diverse, with an abundance of SRM lineages. The evolution of oxygenic photosynthesis has led to the steady and inexorable accumulation of O_2_ in Earth’s atmosphere (the great oxidation event, GOE), with the rise of atmospheric O_2_ to 1-5% of current levels between 2.4 and 2.1 billion years (Gyr) ago, and its accumulation to values comparable to modern values 500-600 Mya ^63^. Due to the sensitivity and expected lack of adaptive mechanisms to cope with atmospheric oxygen in multiple strict anaerobes, as well as the chemical instability of multiple S species in an oxygenated atmosphere, the GOE exerted a profound negative impact on anaerobic surficial life forms (the oxygen catastrophe) leading to the first and arguably most profound extinction event in earth’s history. In addition to suppressing anaerobiosis in atmospherically-exposed habitats, the GOE also led to a significant change in the S cycle, from one based on atmospheric inputs to one dependent on oxidative weathering leading to the release of huge amount of sulfate derived from the oxidation of pyrite and the dissolution of sulfate minerals ^64,^ hitherto a minor byproduct of Archean abiotic and biotic reactions ^15, 65^. Therefore, it appears that loss of niches associated with geological transformations could be one of the possible explanations for high extinction rates for microorganisms on earth, as well as the constant identification of rare, novel taxa within anaerobic settings. It is notable that phylogenetically novel branches with extremely rare distribution in earth (defined as phyla with 5 genomes or less in GTDB) have been consistently identified in anaerobic habitats.

The hypothesis that the GOE played an important role in the extinction of many novel anaerobic lineages, specifically those mediating SCI reductive processes is bolstered by our analysis of the overlaying microbial community in the spring. Here, spatial oxygen intrusion provides a mimicking effect to the relentless temporal progression of oxygenation through the Proterozoic (2.4-2.1 Gya) and Paleo-Phanerozoic (500-600 Mya). The analysis demonstrates a community transitioning from ancient novelty to modern commonality, with the proportion of novel and rare taxa as well as lineages associated with reductive sulfur processes decreasing at the expense of propagation of organisms mediating oxidative sulfur processes.

In summary, by examining microbial diversity in Zodletone spring, we greatly expand the overall diversity within the tree of life via the discovery and characterization of a wide range of novel lineages, and significantly enrich representation of a wide range of LRD lineages. We also describe a unique sulfur-cycling community in the spring that is largely dependent on sulfite, thiosulfate, sulfur, and tetrathionate, rather than sulfate, as an electron acceptor. Given the remarkable similarity to conditions prevailing prior to the GOE, we consider the spring an invaluable portal to investigate the community thriving on the earth’s surface during these eons, and posit that GOE-precipitated the near extinction of a wide range of phylogenetically distinct oxygen-sensitive lineages and drastically altered the reductive sulfur-cycling community from sulfite, sulfur, and thiosulfate reducers to predominantly sulfate reduction in the current earth.

## Materials and Methods

### Site description and geochemistry

Zodletone spring is located in the Anadarko Basin of western Oklahoma (N34.99562° W98.68895°). The spring arises from underground, where water is pumped out slowly along with sediments. Sediments settled at the source of the spring, a boxed square 1m^2^ (Figure S1) are overlaid with water that collects and settles in a concrete pool erected in the early 1900s. The settled water is 50-cm deep above the sediments and is exposed to atmospheric air. Water and sediments originating from the spring source are highly reduced due to the high dissolved sulfide levels (8-10 mM) in the spring sediments. Microsensor measurements show a completely anoxic (oxygen levels < 0.1 μM) and highly reduced source sediments. Oxygen levels slowly increase in the overlaid water column from 2–4 μM at the 2 mm above the source to complete oxygen exposure on the top of the water column ^66^. The spring geochemistry has regularly been monitored during the last two decades ^66, 67, 68^ and is remarkably stable. The spring is characterized by low levels of sulfate (50-94 μM), with higher levels of sulfite (0.21 mM), elemental sulfur (0.1 mM), and thiosulfate (0.52) ^68, 69^.

### Sampling

Samples were collected from the source sediments and standing overlaid water in sterile containers and kept on ice until brought back to the lab (~2h drive), where they were immediately processed. For metatranscriptomics, samples were collected at three different time points: morning (9:15 am), afternoon (2:30 pm), and evening (5:30 pm) in June 2019; stored on dry ice until transferred to the lab where they were stored at −80°C until processed for RNA extraction within a week.

### Nucleic acid extraction

DNA was directly extracted from 0.5 grams of source sediments. For water samples, water was filtered on 0.2 μm sterile filters. DNA was directly extracted from filters (20 filters, 10 L of water samples). Extraction was conducted using the DNeasy PowerSoil kit (Qiagen, Valencia, CA, USA). RNA was extracted from 0.5 g sediment samples using RNeasy PowerSoil Total RNA Kit (Qiagen, Valencia, CA, USA) according to the manufacturer’s instructions.

### 16S rRNA gene amplification, sequencing, and analysis

Triplicate DNA extractions were performed for both sediment and water samples from the Zoddletone spring. To characterize the microbial diversity based on 16S rRNA gene sequences we used the Quick-16S™ NGS Library Prep Kit (Zymo Research, Irvine CA), following the manufacturer’s protocol. For amplification of the V4 hypervariable region we used a mix of modified versions of primers 515F-806R ^70,^ tailored to provide better coverage for several under-represented microbial lineages. They included 515FY (5’GTGYCAGCMGCCGCGGTAA) ^71,^ 515F-Cren (5’GTGKCAGCMGCCGCGGTAA, for Crenarchaeota) ^72,^ 515F-Nano (5’GTGGCAGYCGCCRCGGKAA, for Nanoarchaeota) ^72,^ 515F-TM7 (5’GTGCCAGCMGCCGCGGTCA for TM7/Saccharibacteria) ^73^ as forward mix and 805RB (5’ GGACTACNVGGGTWTCTAAT) ^74^ and 805R-Nano (5’GGAMTACHGGGGTCTCTAAT, for Nanoarchaeota) ^72^ as reverse mix. Purified barcoded amplicon libraries were sequenced on an Illumina MiSeq instrument (Illumina Inc., San Diego, CA) using a v2 500 cycle kit, according to manufacturer’s protocol. Demultiplexed forward and reverse reads were imported as paired fastq files into QIIME2 v. 2020.8 ^75^ for analysis. The DADA2 plugin was used to trim, denoise, pair, purge chimeras and select amplicon sequence variants (ASVs), using the command “qiime dada2 denoise-paired”. Between 44k and 194k non-chimeric sequences were obtained for the individual samples. The ASVs were taxonomically classified in QIIME2 using a trained classifier built based on Silva-138-99 rRNA sequence database. The ASVs were assigned to 1643 taxonomic categories corresponding to taxonomic level 7 (species and above) and to 932 genera (level 6). Alpha rarefaction curves indicated saturation of observed sequence features (ASVs) at a sequencing depth of 70-80k sequences.

### Metagenome sequencing, assembly, and binning

Metagenomic sequencing was conducted using the services of a commercial provider (Novogene, Beijing, China) using two lanes of the Illumina HiSeq 2500 system for each of the water and sediment samples. Transcriptomic sequencing using Illumina HiSeq 2500 2□×□150bp paired-end technology was conducted using the services of a commercial provider (Novogene Corporation, Beijing, China). Metagenomic reads were assessed for quality using FastQC followed by quality filtering and trimming using Trimmomatic v0.38 ^76^. High quality reads were assembled into contigs using MegaHit (v.1.1.3) with minimum Kmer of 27, maximum kmer of 127, Kmer step of 10, and minimum contig length of 1000 bp. Bowtie2 was used to calculate sequencing coverage of each contig by mapping the raw reads back to the contigs. Assembled contigs were searched for ribosomal protein S3 (rpS3) sequences using a custom hidden Markov model (HMM) built from Uniprot reference sequences assigned to the Kegg Orthologies K02982, and K02984 (corresponding to the bacterial, and archaeal RPS3, respectively) using hmmbuild (HMMER 3.1b2). rpS3 Sequences were clustered at 99% ID using CD-HIT as previously suggested for a putative species cutoff for rpS3 data ^77^. Taxonomic affiliations of (rpS3) groups were identified using Diamond Blast against the GTDB r95 database ^19^.

Contigs from the sediment and water assemblies were binned into draft genomes using both Metabat ^78^ and MaxBin2 ^79^. DasTool was used to select the highest quality bins from each metagenome assembly ^80^. CheckM was used for estimation of genome completeness, strain heterogeneity, and contamination ^81^. Genomic bins showing contamination levels higher than 10%, were further refined based on the taxonomic affiliations of the binned contigs, as well as the GC content, tetranucleotide frequency, and coverage levels using RefineM ^82^. Low quality bins (>10% contamination) were cleaned by removal of the identified outlier contigs, and the percentage completeness and contamination were again re-checked using CheckM.

### Genomes classification, annotation, and metabolic analysis

Taxonomic classifications followed the Genome Taxonomy Database (GTDB) release r95 ^19,^ and were carried out using the classify_workflow in GTDB-Tk (v1.1.0) ^83^. Phylogenomic analysis utilized the concatenated alignment of a set of 120 single-copy bacterial genes, and 122 single-copy archaeal genes ^19^ generated by the GTDB-Tk. Maximum-likelihood phylogenomic tree was constructed in FastTree using the default parameters ^84^.

### Annotation and metabolic analysis

Protein-coding genes were predicted using Prodigal ^85^. GhostKOALA ^86^ was used for the functional annotation of every predicted open reading frame in every genomic bin and to assign protein-coding genes to KEGG orthologies (KOs).

### Analysis of sulfur cycling genes

To identify taxa mediating key sulfur-transformation processes in the spring sediments, we mapped the distribution of key sulfur-cycling genes in all genomes and deduced capacities in individual genomes by documenting the occurrence of entire pathways (as explained below in details). This was subsequently confirmed by phylogenetic analysis and examining contiguous genes organization in processes requiring multi-subunit and/or multi-gene. Further, expression data was used from three time points to identify the fraction of the community that is metabolically actively involved in the process. Analysis of S cycling capabilities was conducted on individual genomic bins by building and scanning hidden markov model (HMM) profiles as explained below. To build the sulfur-genes HMM profiles, Uniprot reference sequences for all genes with an assigned KO number were downloaded, aligned using Clustal-omega ^87,^ and the alignment was used to build an HMM profile using hmmbuild (HMMER 3.1b2) ^88^. For genes not assigned a KO number (e.g. *otr*, *tsdA*, *tetH*), a representative protein was compared against the KEGG Genes database using Blastp and significant hits (those with e-values < e-80) were downloaded and used to build HMM profiles as explained above. The custom-built HMM profiles were then used to scan the analyzed genomes for significant hits using hmmscan (HMMER 3.1b2) ^88^ with the option −T 100 to limit the results to only those profiles with an alignment score of at least 100. Further confirmation was achieved through phylogenetic assessment and tree building procedures, in which potential candidates identified by hmmscan were aligned to the reference sequences used to build the custom HMM profiles using Clustal-omega ^87,^ followed by maximum likelihood phylogenetic tree construction using FastTree ^84^. Only candidates clustering with reference sequences were deemed true hits and were assigned to the corresponding KO. Details on the genes examined for evidence of sulfate, sulfite, polysulfide, tetrathionate, and thiosulfate reduction, thiosulfate disproportionation, and various sulfur oxidation processes are provided in the supplementary document (S. text).

#### Phylogenetic analysis and operon organization of S cycling genes

The phylogenetic affiliation of the S cycling proteins AsrB, Otr, PhsC, PsrC, and DsrAB was examined by aligning Zodletone genome predicted protein sequences to Uniprot reference sequences using Mafft ^89^. The DsrA and DsrB alignments were concatenated in MEGA X ^90^. All alignments were used to construct maximum likelihood phylogenetic trees in RAxML ^91^. The R package genoPlotR ^92^ was used to produce gene maps for the DSR and ASR loci in Zodletone genomes using the Prodigal predicted gene starts, ends, and strand.

#### Transcription of sulfur cycling genes

A total of 21.4 M, 27.9 M, and 22.5 M 150-bp paired-end reads were obtained from the morning, afternoon, and evening RNA-seq libraries. Reads were pseudo-aligned to all Prodigal-predicted genes from all genomes using Kallisto with default settings ^93^. The calculated transcripts per million (TPM) were used to obtain total transcription levels for genes identified from genomic analysis as involved in S cycling in the spring.

### Additional metabolic analysis

For all other non-sulfur related functional predictions, combined GhostKOALA outputs of all genomes belonging to a certain order (for orders with 5 genomes or less; n=206), or family (for orders with more than 5 genomes; n=85) were checked for the presence of groups of KOs constituting metabolic pathways (additional file 1). The list of these 291 lineages is shown in Table S2. The presence of at least 80% of KOs assigned to a certain pathway in at least one genome belonging to a certain order/family was used as an indication of the presence of that pathway in that order/family. Such criteria were used for the prediction of autotrophic capabilities, as well as catabolic heterotrophic degradation capabilities of sugars, amino acids, long-chain fatty acids, short chain fatty acids, anaerobic benzoate degradation, anaerobic short chain alkane degradation, aerobic respiration, nitrate reduction, nitrification, and chlorophyll biosynthesis. Glycolytic, and fermentation capabilities were predicted by feeding the GhostKOALA output to KeggDecoder ^94^. Proteases, peptidases, and protease inhibitors were identified using Blastp against the Merops database ^95,^ while CAZymes (glycoside hydrolases [GHs], polysaccharide lyases [PLs], and carbohydrate esterases [CEs]) were identified by searching all ORFs from all genomes against the dbCAN hidden Markov models V9 ^96^ (downloaded from the dbCAN web server in September 2020) using hmmscan. FeGenie ^97^ was used to predict the presence of iron reduction and iron oxidation genes in individual bins.

## Acknowledgments

This work was supported by the National Science Foundation Grants 2016423 (to NHY and MSE) and 2016371 to MP.

