## Supplementary document for "Microbial diversity and sulfur cycling in an early earth analogue: From ancient novelty to modern commonality"

**Supplementary Text**

**I. Methods**

**Analysis of sulfur cycling genes.**

*Sulfate-reduction.* Sulfate reduction capacity was assessed by the presence of genes encoding the enzymes 3'-phosphoadenosine 5'-phosphosulfate synthase [Sat; EC:2.7.7.4 2.7.1.25] for sulfate activation to adenylyl sulfate (APS), the enzyme complex adenylylsulfate reductase [AprAB; EC:1.8.99.2] for APS reduction to sulfite, the quinone-interacting membrane-bound oxidoreductase complex [QmoABC] for electron transfer, the enzyme dissimilatory sulfite reductase [DsrAB; EC:1.8.99.5] and its co-substrate DsrC for dissimilatory sulfite reduction to sulfide, and the sulfite reduction-associated membrane complex DsrMKJOP for linking cytoplasmic sulfite reduction to energy conservation.

*Sulfite-reduction.* Sulfite could be utilized by most sulfate-reducing microorganisms ^1^. Dedicated sulfite-reduction capacity was assessed by the presence of the dissimilatory sulfite reductase system explained above ^2, 3^ with the lack of sulfate-activation (Sat) and reduction (Apr) genes. In addition, sulfite-reduction was assessed via the sole or co-occurrence of the anaerobic sulfite reductase (AsrABC) system ^4^, along with the membrane-bound associated complex (HdrABC) for transfer of electrons to the AsrC subunit ^5^. The Asr enzyme has been shown to function in the cytoplasm in *Salmonella typhimurium* to reduce the sulfite released from respiratory reduction of tetrathionate and thiosulfate ^4^. However, a scenario where the Asr enzyme is involved in sulfite respiration is possible via electron transfer from a membrane-bound associated complex to AsrC (the physiological partner of AsrAB). A plausible candidate for this membrane complex is the heterodisulfide reductase-related enzymes (HdrABC), analogous to what was suggested for DsrC (the physiological partner of DsrAB) in organisms lacking the sulfite reduction-associated membrane complex DsrMKJOP ^5^.

*Polysulfide reduction:* In addition to sulfate and sulfite, Zodletone spring is euxinic with extremely high levels of zero valent sulfur, available as soluble polysulfide. Respiratory polysulfide reduction was assessed via the identification of the membrane-bound molybdoenzyme complex PsrABC, which reduces polysulfides with electrons obtained from either a hydrogenase or a formate dehydrogenase through a quinone electron carrier ^6^. In addition to the membrane-bound Psr system, representatives of the cytolpasmic sulfurhydrogenase I (HydABCD system), and/or II (ShyABCD system) were identified. However, although these enzymes have been shown to be dissimilatory in the archaeon *Pyrococcus furiosus* ^7, 8^, their involvement in an ETS-associated respiration is currently unclear.

*Thiosulfate reduction/ disproportionation:* Thiosulfate occurs in natural environments as a result of the reaction of sulfite with bisulfide (HS^-^) ^9^. Thiosulfate is relatively stable at neutral pH and is present in high levels in Zodletone spring, Thiosulfate contains two sulfur atoms: a sulfone-sulfur (oxidation state +5), and a sulfane-sulfur (oxidation state -1). As such, thiosulfate can be disproportionated where the sulfone-sulfur is reduced (serves as an electron acceptor), and the sulfane-sulfur is oxidized (serves as an electron donor), with the products being hydrogen sulfide, and sulfite, respectively. We searched for genes encoding the three known pathways for thiosulfate-disproportionation. First, in pure cultures of several sulfate reducers in the Desulfobacterota and Firmicutes, e.g. *Desulfovibrio*, *Desulfotomaculum*, thiosulfate disproportionation is known to occur via a cytochrome c-dependent thiosulfate reductase [EC: 1.8.2.5] ^10, 11, 12, 13, 14, 15, 16, 17^. Second, in pure culture members of the family Enterobacteriaceae (Gammaproteobacteria), thiosulfate disproportionation is known to occur via the quinone-dependent membrane-bound molybdopterin-containing thiosulfate reductase PhsABC ^18^. Finally, thiosulfate disproportionation to sulfite and hydrogen sulfide can also occur via a rhodanase-like enzyme [EC: 2.8.1.1 or EC: 2.8.1.3], as shown for several bacterial lineages ^19, 20, 21, 22, 23^, although this could be part of a thiosulfate assimilatory pathway as recently shown in *E. coli* ^24^.

Following the disproportionation of thiosulfate to sulfite and hydrogen sulfide, microorganisms differ in the fate of the produced sulfite. Some microorganisms reduce the released sulfite to sulfide via a Dsr or Asr dissimilatory sulfite reductase ^18^), leading to complete reduction of one thiosulfate molecule to two sulfides (thiosulfate-reduction). Others oxidize the released sulfite to sulfate via the reversal of the sulfate reduction pathway ^17, 25^, or via the sulfite dehydrogenases SorAB or SoeABC ^26^, leading to the final conversion of one thiosulfate molecule to one sulfide and one sulfate molecules. The distribution of all thiosulfate disproportionation capacities were assessed by the occurrence of one of the three pathways described above, and the fate of sulfite in genomes mediating the initial disproportionation steps was assessed as described above.

*Tetrathionate reduction:* Tetrathionate has two sulfur atoms in oxidation state of 0 while the other two are in oxidation state of +5. In nature, tetrathionate is formed via the biotic or abiotic oxidation of thiosulfate under anoxic conditions ^9^. Some microorganisms are capable of tetrathionate respiration via membrane-bound tetrathionate reductases that will reduce tetrathionate to thiosulfate serving as the terminal oxidase in a short electron transport system. Enzymes mediating such process include octaheme tetrathionate reductase (otr) ^27^, as well as the guanylyl molybdenum cofactor-containing tetrathionate reductase (ttrABC) ^28^. The produced thiosulfate could be metabolized through disproportionation as described above.

*Oxidative sulfur processes.* The versatile sulfur oxidation (SOX enzyme complex) system was assessed in all genomes. The SOX system mediates the oxidation of a wide range of reduced sulfur compounds (sulfide, sulfite, thiosulfate, and elemental sulfur) directly to sulfate*.* Sulfide oxidation to sulfur was also assessed by the presence of the sulfide dehydrogenase FccAB [EC: 1.8.2.3] and/or the sulfide:quinone oxidoreductase Sqr [EC: 1.8.5.4], both known to oxidize sulfide to sulfur or polysulfide. Sulfur/polysulfide oxidation to sulfite was assessed via the reversal of the Dsr system (encompassing the full Dsr system *dsrAB*+*dsrC*+*dsrMKJOP*, in addition to the genes *dsrEFH*, *tusA*, and *rhdA*). Sulfite oxidation to sulfate was assessed via the reversal of AprAB+QmoABC system, the sulfite dehydrogenase (quinone) SoeABC [EC: 1.8.5.6], or the sulfite dehydrogenase (c-type cytochrome) SorAB [EC: 1.8.2.1]. Thiosulfate oxidation to tetrathionate was assessed via the thiosulfate dehydrogenase *tsdA* [EC: 1.8.2.2], or the thiosulfate dehydrogenase (quinone) *doxAD* [EC: 1.8.5.2]. Tetrathionate generated could be cleaved using tetrathionate hydrolase (*tetH*) ^29^ that is known to cleave tetrathionate to thiosulfate, sulfur, and sulfate, or converted to sulfite using the rDSR system.

**II. Results.**

**1. Detailed phylogenomic analysis of Zodletone spring sediments and water communities.**

***Anoxic sediments.***

*Overall sequencing overview and diversity patterns.* Metagenomic sequencing of the spring sediments yielded 281 Gbp, 79.54% of which assembled into 12 Gbp contigs, with 6.8 Gbp contigs longer than 1Kbp. 1,848 genomes were binned, 683 of which passed quality control criteria, and 516 remained after dereplication (Table S1). These MAGs represented 64 phyla or candidate phyla (53 bacterial and 11 archaeal), 127 classes, 198 orders, and 300 families (Figure 1a-b). Diversity assessment utilizing small subunit ribosomal protein S3 from assembled contigs (n=2079), as well as a complementary 16S rRNA illumina sequencing effort (n=309,074 amplicons), identified a higher number of taxa (82 phyla and 1679 species in the ribosomal protein S3 dataset, and 69 phyla and 1050 species in 16S rRNA dataset) (Figure S2). Nevertheless, the overall community composition profiles generated from all three approaches were broadly similar.

*Detailed phylogenetic analysis of Zodletone sediment MAGs.* The Chloroflexota (n=69), Planctomycetota (n=47), Bacteroidota (n= 43), Desulfobacterota (n= 43), Spirochaetota (n= 28 genomes), Patescibacteria (n=20 genomes), and the archaeal phylum Nanoarchaeota (n=21) were the most abundant phyla in Zodletone spring sediments, albeit representing only 52.52% of the total number of recovered genomes (Figure 1, 2C). A notable absence, or extreme paucity, of genomes belonging to the Proteobacteria (6 genomes) and Firmicutes (12 genomes), the most successful taxa within current biomes ^30^, as well as the oxygen-generating oxycayanobacteria (0 genomes) were observed (Figure 1, 2C). Within the Chloroflexota, 38/69 genomes belonged to 3 novel orders, 5 novel families, and multiple LRD orders (Thermoflexales, 4572-78, and UBA2777) and families (E44-bin32, Fen-1058, J111, RBG-13-53-26, RBG-16-64-43, UBA11579, UBA11858, UBA2029, UBA2162, UBA3940, UBA4811, UBA4823, UBA5620, UBA5760, and UBA6092) (Figure S3a). Within the Planctomycetota, 17/47 genomes belonged to 2 novel orders, 8 novel families, and multiple LRD orders (FEN-1346, SZUA-567, and UBA8890) and families (Fen-1342, SM23-30, UBA1845, UTPLA1, and UBA8108, Figure S3b). Within the Bacteroidota, 27/43 genomes belonged to 1 novel family, and multiple LRD families (FEN-979, GCA-2748055, NBLH01, UBA10428, UBA12170, UBA5072, UBA6680, and SZUA-365, Figure S3c). Within the Spirochaetota, 19/28 genomes belonged to one novel class, 2 novel orders, and 9 novel families, as well as multiple LRD families (ARS1246, Marispirochaetaceae, and RPPD01, Figure S3d). Within the Desulfobacterota (Figure S3e), 35/43 genomes belonging to 3 novel classes, 10 novel orders, and 7 novel families were identified, as well as multiple LRD families (B25-G16, BuS5, HGW-15, MLS-D, NaphS2, UBA2210, and UBA3084). Only 6/43 genomes belonged to the well-described families Desulfovibrionaceae, Geopsychrobacteraceae, Smithellaceae, and Syntrophaceae. Finally, an extremely diverse community of Patescibacteria (13 different orders, 3 of which belonging to LRD orders, and 14 different families, including 6 novel and 2 LRD families), and Nanoarchaeota (2 orders including the LRD order CG07-land), and 15 different families, including 5 novel and 10 LRD families) were identified in the spring sediments (Figure S3F). A similar pattern of high proportion of novel and LRD families was identified throughout all other lineages (Figure 1a). Therefore, in addition to expanding the number of novel lineages (classes, orders, and families), and greatly enriching available genomes in rare, poorly represented taxa, our results highlight the uniqueness and distinction of the microbial community thriving in Zodletone spring sediments, compared to all previously studied habitats on the current earth.

***Hypoxic waters*.**

*Overall sequencing overview and diversity patterns.* Metagenomic sequencing of the oxygen-exposed overlaying water column community yielded 323 Gbp, 80.07% of which assembled into 3.6 Gbp contigs with 3.1 Gbp contigs >1K. 883 genomes were binned, with only 114 remaining after dereplication. Of these, 62 belonged to shared families with the sediment community, and 52 were water specific.

*Detailed phylogenetic analysis of Zodletone water MAGs.* Water-specific genomes (n=52) mostly belonged to well-characterized microbial lineages, e.g. class Alphaproteobacteria (5 genomes belonging to Rhodobacteraceae and Rhodospirillaceae, and 3 belonging to the uncultured lineages NBLK01 and Rs-D84), Gammaproteobacteria (9 genomes belonging to the lineages Thiomicrospiraceae, Halothiobacillaceae, Acidithiobacillaceae, Burkholderiaceae, Chromatiaceae, and Methylothermaceae, and 1 belonging to the uncultured lineage UBA9339), phylum Camplylobacterota (8 genomes, belonging to the families Sulfurimonadaceae, Sulfurovaceae, and Sulfurospirillaceae), Firmicutes/Firmicutes_A (4 genomes, and 6 genomes belonging to the Bacilli, and Clostridia classes, respectively), and well described families in the phyla Bacteroidota (Flavobacteriaceae, Prolixibacteraceae, Paludibacteraceae, Marinilabiliaceae, Tannerellaceae, Marinifilaceae, Balneolaceae), Desulfobacterota (Families Geopsychrobacteraceae, Desulfuromonadaceae,), and Spriochaetota (Sphaerochaetaceae, Treponemataceae, Spirochaetaceae_B). Collectively, this demonstrates a pattern where the intrusion of oxygen is associated with a negative impact on novel and LRD lineages that are prevalent in the sediment, and the propagation of communities associated with well-characterized lineages within the bacterial tree of life.

**3. Transcriptomic analysis.** Transcriptional expression of genes involved in S-species reduction/disproportionation was analyzed, and the identity of the active sulfur-reducing community in the spring sediment was examined (Figure 5). All S-species reduction/ disproportionation genes discussed above were identified in the metatranscriptomic dataset, and transcripts belonging to 51 different phyla were identified. Analysis of the spring sediment revealed the transcription of both the Dsr and Asr systems for sulfite reduction with contributions from Chloroflexota, Planctomycetota, Desulfobacterota, Bacteroidota, Fermentibacterota, Acidobacteriota, CSSED10-310, Actinobacteriota, Spirochaetota, WOR-3, and Fibrobacterota (Asr), and Desulfobacterota, Acidobacteriota, Zixibacteria, and Myxococcota (Dsr). Sulfate reduction genes (Sat, AprAB, and QmoABC) were also transcribed with major contribution from Desulfobacterota, Myxococcota, Zixibacteria, and Acidobacteriota. Total transcription levels of the Asr system were 4-times higher than the Dsr system, consistent with the higher number of Zodletone sediment genomes encoding the Asr system compared to the Dsr system. Transcription of the thiosulfate disproportionating rhodanese-like enzyme [EC: 2.8.1.1 or EC: 2.8.1.3] was detected in the phyla Desulfobacterota, Actinobacteriota, Firmicutes_A, Chloroflexota, Fibrobacterota, Planctomycetota, Spirochaetota, Bacteroidota, Halobacteriota, and Acidobacteriota, while the transcription of the thiosulfate reductase *phsABC* was detected in the phyla Actinobacteriota and Bacteroidota. Transcription of the tetrathionate reduction genes *ttrABC* was detected in the phyla Desulfobacterota, Actinobacteriota, Bacteroidota, Chloroflexota, Acidobacteriota, and Spirochaetota, while the octaheme tetrathionate reductase *otr* transcription was detected in Desulfobacterota, Bacteroidota, Myxococcota, UBP7_A, and Chloroflexota. Finally, the transcription of *psrABC* for polysulfide reduction was detected majorly in the phyla Bacteroidota, Desulfobacterota, and Campylobacterota, while transcription of the cytoplasmic sulfurhydrogenases I and II (*hyd/shy* systems) was identified in the phyla Actinobacteriota, Chloroflexota, Planctomycetota, Myxococcota, Acidobacteriota, Bacteroidota, and Desulfobacterota.

**4.** **Oxidative sulfur processes dominate Zodletone water community.**

In contrast to sediment communities, reductive sulfur-processes were identified in only 25 (21.92%) water genomes, as opposed to 149 (29%) sediment genomes (Figure 3, Table S3). Dissimilatory sulfate reduction to sulfide capacity was completely absent in water genomes. The capacity for dissimilatory sulfite reduction via the Dsr system was absent, and the Asr system was only encoded in 7 water genomes. Thiosulfate reduction/disproportionation capacity to sulfide and sulfate (PhsABC + SoeABC, and/or Rhodanase + Dsr/Asr + SoeABC/SorAB) was encoded in only four genomes, all of which also encoded the capacity for tetrathionate reduction (via Otr and/or ttrABC). Finally, respiratory polysulfide reduction (via PsrABC) was encoded in 19 genomes. In all cases, the reductive sulfur community in water was a subset of the sediment community.

In contrast, oxidative sulfur processes dominated the water community, with pathways encoding sulfide, sulfur, thiosulfate, tetrathionate, and/or sulfite oxidation to sulfate present in 59/114 genomes (51.8% of all water genomes) belonging to 13 phyla, 16 classes, 25 orders, and 43 families. The oxidative sulfur community in the water belonged to mostly well-characterized lineages (Table S3, Figure 3). Only 8 and 10 genomes involved in oxidative sulfur processes belonged to novel, and LDR families, respectively.

A complete SOX system, putatively mediating oxidation of a wide range of reduced sulfur-species to sulfate was encoded in genomes belonging to well-characterized families within the Proteobacteria (11 genomes total belonging to families Acidithiobacillaceae, Burkholderiaceae, Halothiobacillaceae, Rhodobacteraceae, and Thiomicrospiraceae) and Campylobacterota (3 genomes in the family Sulfurimonadaceae).

*Sulfide oxidation to sulfur and sulfite.* Thirty nine water genomes encoded the sulfide dehydrogenase fccAB [EC: 1.8.2.3] and/or the sulfide:quinone oxidoreductase Sqr [EC: 1.8.5.4] both known to oxidize sulfide to sulfur/ polysulfide. These genomes belonged to the phyla Bacteroidota (14 genomes in the well characterized families Chlorobiaceae, Prolixibacteraceae, and Paludibacteracaeae, as well as the uncultured families NBLH01, UBA1556, DTU049, and F082 in the order Bacteriodales), Proteobacteria (13 genomes in the families Acidithiobacillaceae, Burkholderiaceae, Chromatiaceae, Halothiobacillaceae, Methylothermaceae, Rhodobacteraceae, Thiomicrospiraceae), Campylobacterota (8 genomes in the families Sulfurimonadaceae, Sulfurospirillaceae, Sulfurovaceae), in addition to three genomes in the families Anaerolineaceae, Geopsychrobacteraceae, and UBA2242 within the phyla Chloroflexota, Desulfobacterota, and Marinisomatota, respectively, and one genome belonging to a novel Thermodesulfovibrionales family (Nitrospirota). Only two of the above thirty-nine genomes (one Proteobacteria genome and one Nitrospirota genome) encoded the capacity to further oxidize the sulfur/polysulfide to sulfite via the reversal of the Dsr system (encompassing the full Dsr system *dsrAB*+*dsrC*+*dsrMKJOP*, in addition to the genes *dsrEFH*, *tusA*, and *rhdA*).

*Sulfite oxidation to sulfate*: A total of twenty-six water genomes encoded the capacity for sulfite oxidation to sulfate via the reversal of AprAB+QmoABC system (1 Bacteroidales genome), the sulfite dehydrogenase (quinone) SoeABC [EC: 1.8.5.6] (22 genomes belonging to the order Bacteroidales, and the families Acidithiobacillaceae, Burkholderiaceae, Chromatiaceae, Dethiosulfatibacteraceae, Halothiobacillaceae, Methylothermaceae, Rhodobacteraceae, Thiomicrospiraceae within Proteobacteria, the families Sulfurimonadaceae, Sulfurospirillaceae, Sulfurovaceae within Campylobacterota, the Syntrophales family UBA3084, and a novel Thermodesulfovibrionales family (Nitrospirota)), or the sulfite dehydrogenase (cytochrome) SorAB [EC: 1.8.2.1] (3 genomes total within the families Chromatiaceae, Halothiobacillaceae (Proteobacteria), and UBA12059 (Spirochaetota)).

*Thiosulfate oxidation to tetrathionate, and complete thiosulfate oxidation to sulfate via tetrathionate*: Eight water genomes encoded thiosulfate to tetrathionate oxidation capacities via either the thiosulfate dehydrogenase *tsdA* [EC: 1.8.2.2] (7 genomes belonging to the families Sulfurimonadaceae within Campylobacterota, and Rhodobacteraceae, Burkholderiaceae, Halothiobacillaceae, Thiomicrospiraceae within Proteobacteria), or the thiosulfate dehydrogenase (quinone) *doxAD* [EC: 1.8.5.2] (1 Flavobacteriaceae genome). Two of these 8 genomes (1 Rhodobacteraceae, and 1 Halothiobacillaceae genomes) also encoded tetrathionate hydrolase (*tetH*) ^29^ that is known to cleave tetrathionate to thiosulfate, sulfur, and sulfate. Simultaneous identification of the SOX system and both forms of sulfide dehydrogenase (fccAB and Sqr) imply that these two genomes encode the capacity for complete thiosulfate oxidation to sulfate.

*Tetrathionate oxidation*: In addition to the above two genomes, ten other water genomes encoded tetrathionate hydrolase, but with no other means of thiosulfate and sulfide oxidation capacities. Surprisingly, TetH (without other means of thiosulfate or sulfide oxidation) was also encoded in 100 sediment genomes, belonging to 26 phyla, 37 classes, 49 orders, and 66 families (including 23 novel families). Only nine of these genomes (belonging to 9 families including 4 novel ones) showed *tetH* transcriptional levels above 1. However, the exact function of tetrathionate hydrolase in these organisms is not entirely clear, as the subsequent steps of oxidation could not be identified.

**5. Additional metabolic capacities in Zodletone spring sediments.**

In addition to reductive sulfur processes, strict fermentative capacities were highly prevalent in sediment genomes (Table S2), being identified in 100 of the 291 lineages studied. On the other hand, a dearth of aerobic (only 38 lineages), nitrate (only 65 lineages encoded dissimilatory nitrite reduction to ammonium, with 2 of which also encoding the suite of genes for denitrification), Fe^3+^ respiration (8 lineages), or chemolithotrophic nitrifying (only 1 lineage encoded the combination of ammonia monooxygenase and hydroxylamine dehydrogenase), and photosynthetic capacities were identified (Figure 1a, Table S2). Strict fermentative lineages mediate the degradation a wide range of substrates, e.g. sugars (89 of the 100 fermentative lineages), amino acids (85 of the 100 fermentative lineages), short chain fatty acids (37 of the 100 fermentative lineages), complex carbohydrates (36 of the 100 fermentative lineages), long chain fatty acid oxidation (2 lineages), and short chain alkanes (1 lineage) (Table S2), producing a wide range of fermentative end products including lactate, formate, acetate, ethanol, succinate, and hydrogen. Primary productivity in the spring sediments appears to be mostly mediated via hydrogen utilization coupled to either sulfur-cycle intermediates (SCI) reduction (27 lineages, Table S2), or to CO_2_ fixation by hydrogenotrophic methanogens and acetogens using the Wood-Ljungdahl pathway (8 lineages Table S2).

**Supplementary Figures:**

**Figure S1. Zodletone spring source sediments and overlaid water.**

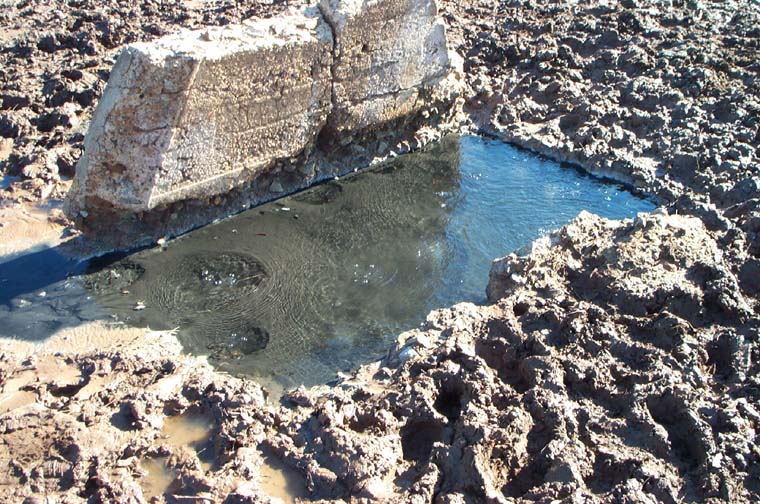

**Figure S2.** Zodletone spring phylum-level community composition based on ribosomal protein S3 (RP-S3), binned genomes (MAGs), as well as the gene for 16S rRNA for both the sediment and the water samples.

Percentage of total

|  | 4572-55 | AABM5-125-24 | Acidobacteriota | Actinobacteriota | Aenigmarchaeota | Aerophobota | Altarchaeota | Armatimonadota | Asgardarchaeota | Aureabacteria | B130-G9 | B64-G9 | Bacteroidota | Bdellovibrionota | Bipolaricaulota | BMS3Abin14 | Caldatribacteriota | Calditrichota | Campylobacterota | CG03 |
| --- | --- | --- | --- | --- | --- | --- | --- | --- | --- | --- | --- | --- | --- | --- | --- | --- | --- | --- | --- | --- |
| Sediment RP-S3 | 1 | 1 | 32 | 48 | 29 | 1 | 14 | 1 | 23 | 5 | 5 | 3 | 121 | 1 | 2 | 1 | 8 | 8 | 4 | 1 |
| Sediment MAGs |  |  | 18 | 16 | 5 |  | 3 | 1 | 6 | 3 | 2 |  | 43 | 1 | 1 | 0 | 1 | 3 | 2 | 1 |
| Sediment 16S rRNA |  | 0.490743 | 1.37751 | 10.7485 | 0.120908 |  | 0.0662729 | 0.00808206 | 0.0307118 |  |  |  | 14.5519 | 0.206578 | 0.0329748 |  | 0.115089 | 0.818228 | 0.201728 |  |
| Water RP-S3 | 0 | 0 | 33 | 80 | 1 | 0 | 3 | 0 | 1 | 0 | 0 | 0 | 298 | 3 | 0 | 1 | 1 | 2 | 82 | 0 |
| Water MAGs |  |  | 0 | 3 | 0 |  | 0 | 0 | 0 | 0 | 0 |  | 29 | 0 | 0 | 1 | 0 | 1 | 8 | 0 |
| Water 16S rRNA |  | 0.369759 | 1.36492 | 8.93429 | 0.0402419 |  | 0.0388811 | 0.00758181 | 0.0136084 |  |  |  | 12.872 | 0.351874 | 0.0118587 |  | 0.114894 | 0.831861 | 6.3137 |  |

**Figure S3.** Phylogenetic affiliation and contig organization of selected sulfur reduction proteins. (A) Phylogeny of the dissimilatory sulfite reductase DsrAB [EC:1.8.99.5] concatenated proteins (A), anaerobic sulfite reductase subunit B AsrB (B), polysulfide reductase subunit gamma PsrC (C), thiosulfate reductase cytochrome b subunit PhsC (D), and octaheme tetrathionate reductase Otr (E). Alignments were created in Mafft ^37^ and maximum likelihood trees were constructed in RaxML ^38^. Bootstrap support values are shown as bubbles for nodes with >50% support. Branches and branch labels are color coded by phylum for Zodletone sequences. Branch labels depict classification to family level followed by the NCBI genome accession number. Reference sequences are shown in black with the Uniprot accession numbers. Contig organizations of the DSR and ASR loci in selected Zodletone genomes are shown to the right of the trees in A and B. Genes are color coded as shown in the top right corner. Unrelated genes are shown by grey arrows. Gene maps were created in R using the package genoplotR ^39^. Phylum/class classification is depicted to the right of the trees in C-E.

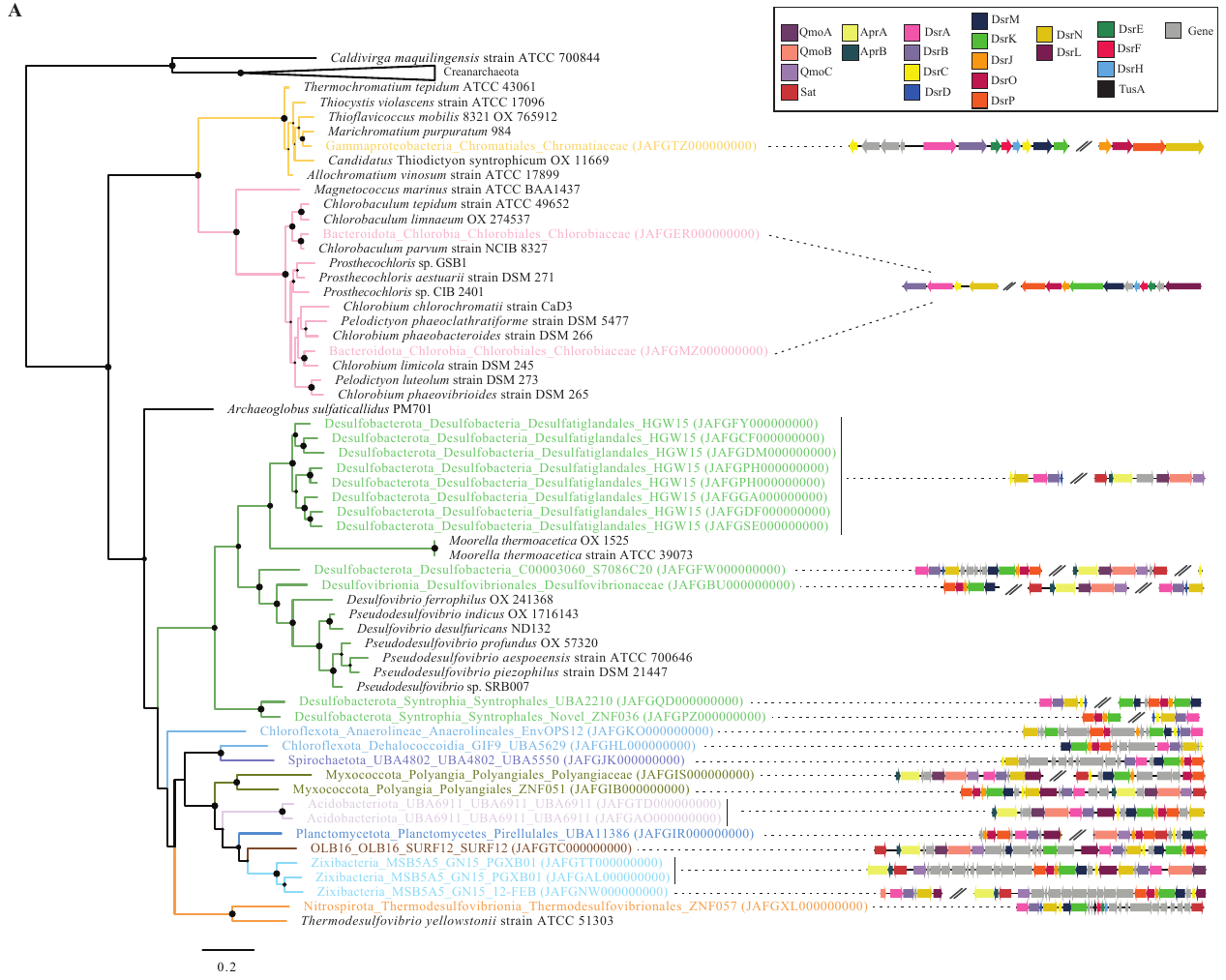

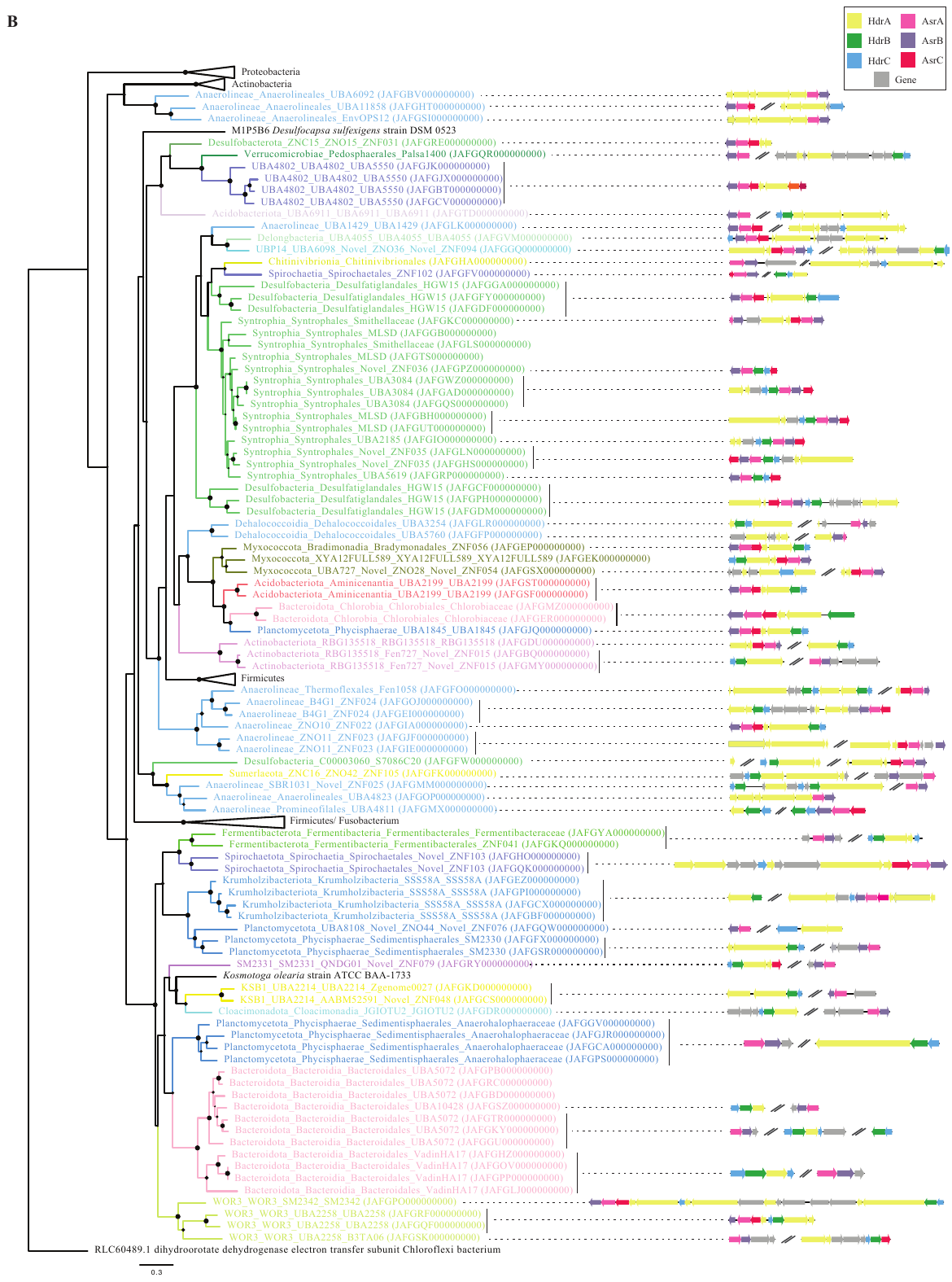

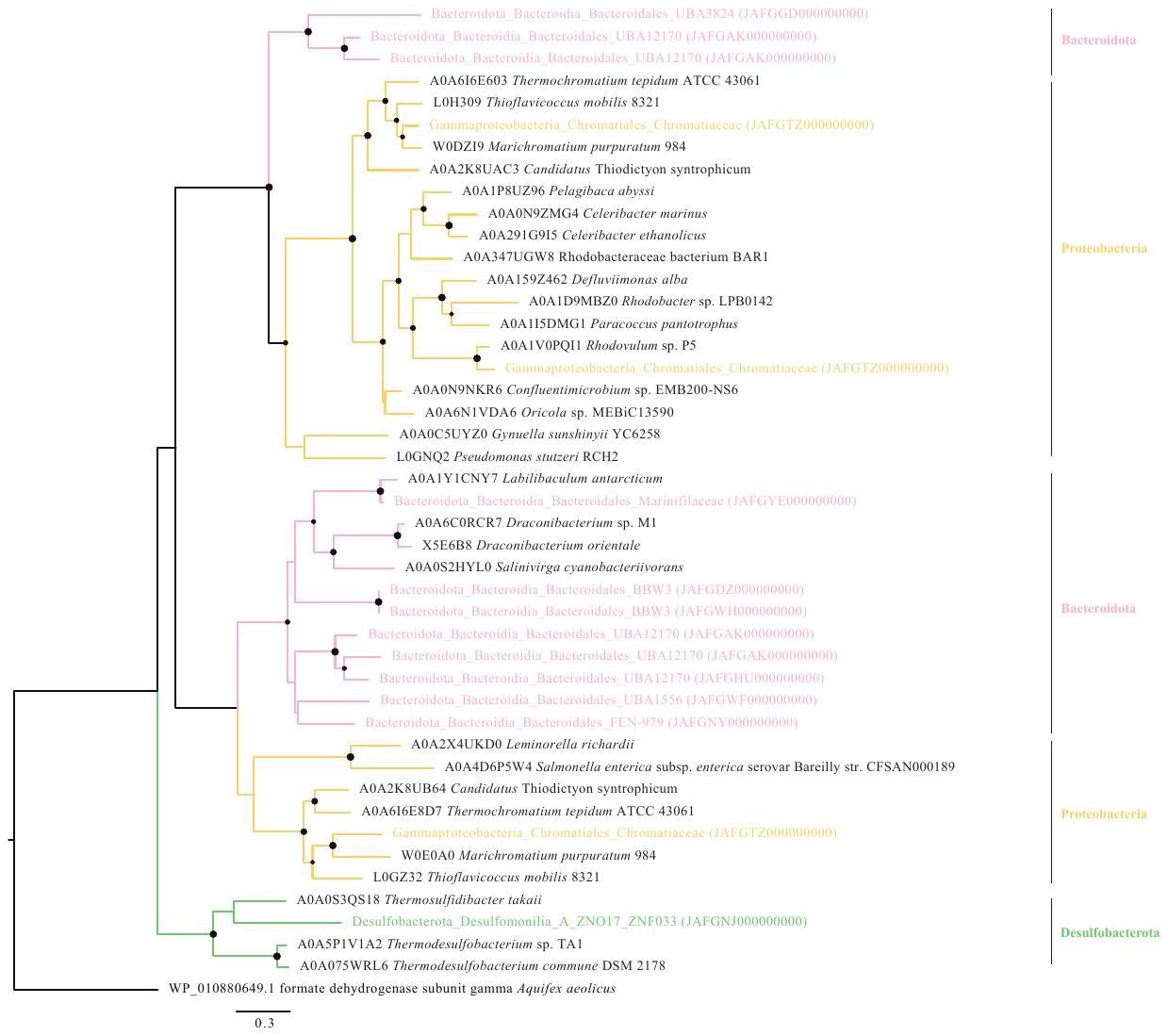

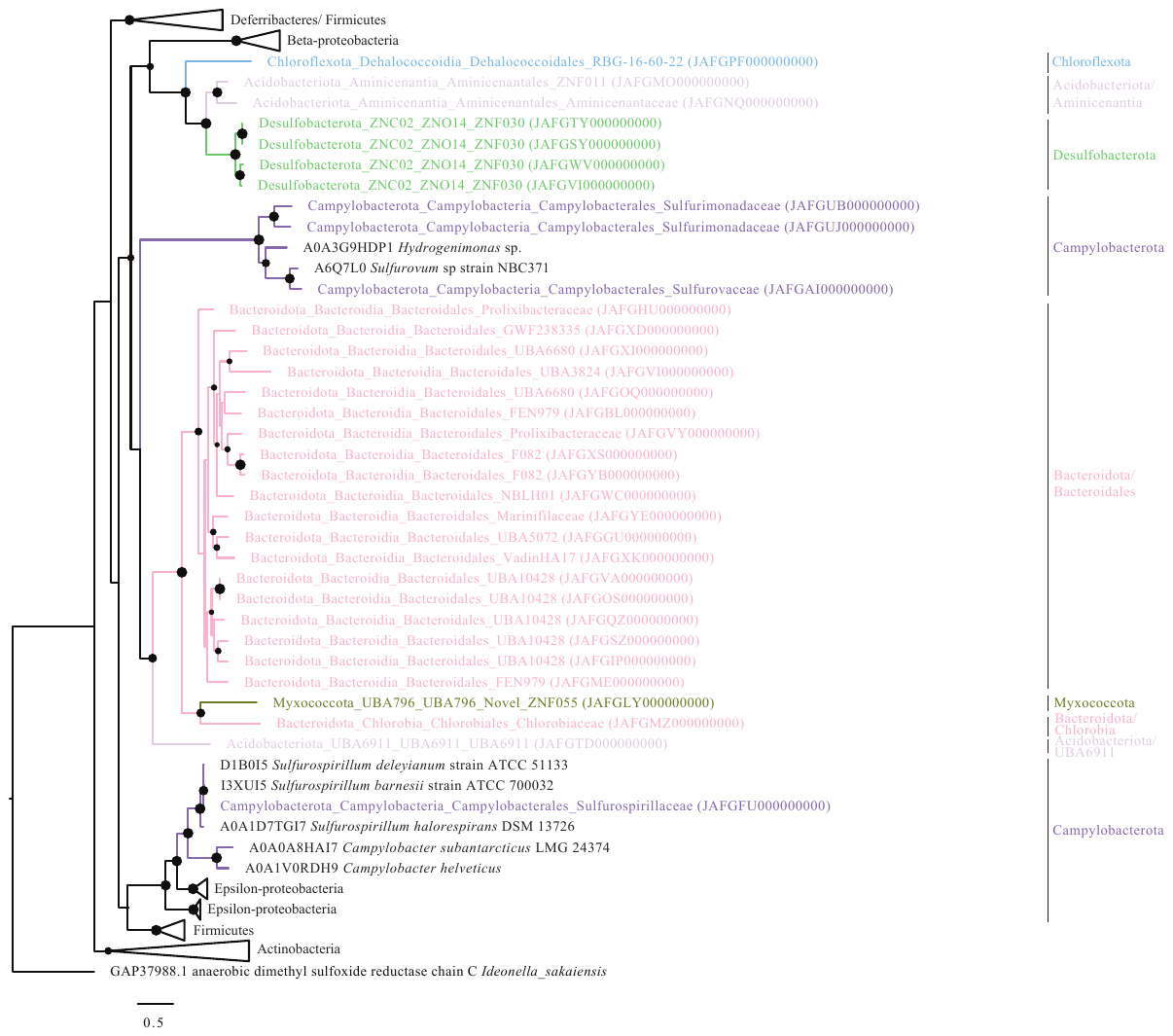

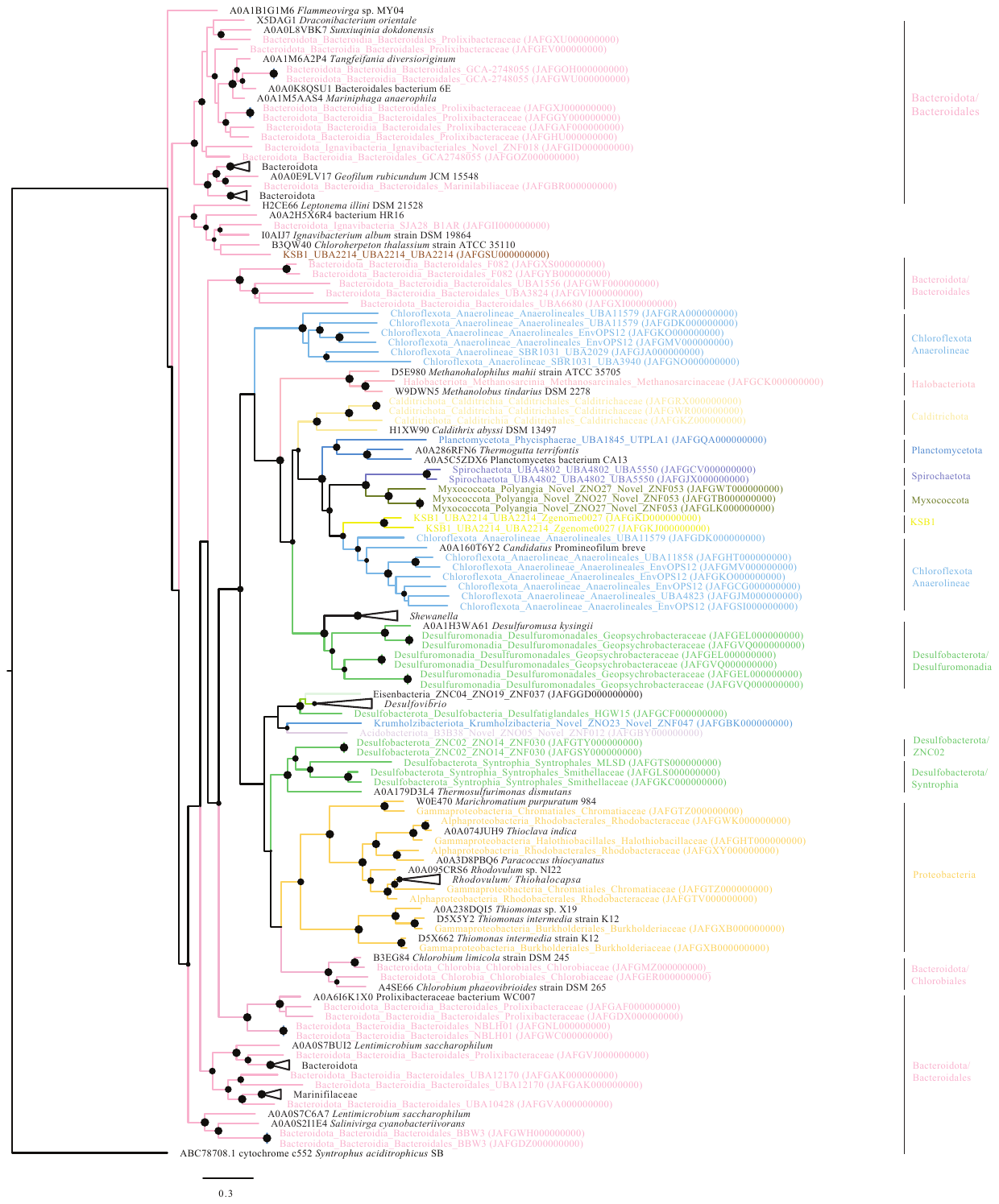

**Supplementary tables:**

**Table S1:** List of all genomes analyzed in this study with their NCBI Assembly accession numbers, taxonomic classification, sequencing statistics, and general genomic features.

**Table S2:** Substrates potentially supporting growth, predicted fermentation end products, and energy conservation pathways predicted from genomic analysis.

**Table S3:** S-cycling genes predicted in Zodletone genomes. Genes are shown in the table header and actual gene names are shown in the corresponding cells.
